# *In situ* structural analysis of the flagellum attachment zone in *Trypanosoma brucei* using cryo-scanning transmission electron tomography

**DOI:** 10.1101/2020.02.14.949115

**Authors:** Sylvain Trépout

**Affiliations:** Institut Curie, Inserm US43, CNRS UMS2016, Université Paris-Sud, Université Paris-Saclay, Centre Universitaire, Bât. 101B-110-111-112, Rue Henri Becquerel, CS 90030, 91401 ORSAY Cedex, FRANCE

**Keywords:** Electron cryo-scanning transmission electron tomography, trypanosome, bloodstream forms, flagellum, flagellum attachment zone (FAZ), FAZ filament

## Abstract

The flagellum of *Trypanosoma brucei* is a 20 µm-long organelle responsible for locomotion and cell morphogenesis. The flagellum attachment zone (FAZ) is a multi-protein complex whose function is to attach the flagellum to the cell body but also to guide cytokinesis. Cryo-transmission electron microscopy is a tool of choice to access the structure of the FAZ in a close-to-native state. However, because of the large dimension of the cell body, the whole FAZ cannot be structurally studied *in situ* at high resolution in 3D using classical transmission electron microscopy approaches. In the present work, cryo-scanning transmission electron tomography, a new method capable of investigating cryo-fixed thick biological samples, has been used to study whole *T. brucei* cells at the bloodstream stage. The method has been used to visualise and characterise the structure and organisation of the FAZ filament. It is composed of an array of cytoplasmic stick-like structures. These sticks are heterogeneously distributed between the posterior part and the anterior tip of the cell. This cryo-STET investigation provides new insights in the structure the FAZ filament. In combination with protein structure predictions, this work proposes a new model for the elongation of the FAZ.

**Highlights:**

- Flagellar and cellular membranes are in close contact next to the FAZ filament
- Sticks are heterogeneously distributed along the FAZ filament length
- Thin appendages are present between the FAZ filament sticks to neighbouring microtubules
- FAZ elongation could originate from the force exerted by dynein motors on subpellicular microtubules

## Introduction

*Trypanosoma brucei* is a unicellular parasite responsible for human African trypanosomiasis, also known as sleeping sickness, occurring in sub-Saharan Africa (Büscher et al., 2017; Rotureau and Van Den Abbeele, 2013). This organism adopts different stages whose shape, intracellular organisation and metabolism vary during the complex life cycle in the insect vector or the mammalian host (bloodstream forms). Reverse genetic approaches such as RNA interference (Ngô et al., 1998), *in situ* tagging (Dean et al., 2015) and more recently CRISPR-Cas9 (Beneke et al., 2017) technologies are potent genetic tools to study gene function of fully sequenced *T. brucei* genome (Berriman et al., 2005; Sistrom et al., 2014). Furthermore, it has a single flagellum during the cell cycle except during cell duplication where a new flagellum (*i.e*. the one of the future daughter cell) is built next to the existing one (Lacomble et al., 2010, 2009; Sherwin and Gull, 1989). Mature cells have a single fully-grown 20 µm-long flagellum. The presence of a single flagellum is an advantage for the study of proteins present in the flagellum, making the phenotype of the inducible mutant cells more easily visible and distinctive than in multiflagellated cells (Blisnick et al., 2014). In *T. brucei*, the flagellum is responsible for cell locomotion (Bastin et al., 1998) and morphogenesis (Kohl et al., 2003). Mechanistically, it has been proposed that the bi-helical swimming pattern of *T. brucei* originates from flagellum motility which is transmitted to the cell body through a succession of structural connecting elements (Heddergott et al., 2012). The sliding model explaining flagellum motility has been first proposed by Peter Satir in 1968 (Satir, 1968). Since then, high resolution cryo-transmission electron microscopy (TEM) revealed that it originates from the force exerted by the outer and inner dynein arms on the 9 microtubules doublets of the axoneme (Lin et al., 2014; Lin and Nicastro, 2018). In the case of trypanosomes, the movement of the axoneme is then transmitted to the paraflagellar rod (PFR), a semi-crystalline multiprotein complex which is a unique feature of most species of the Kinetoplastid order among other eukaryotes (Koyfman et al., 2011; Vickerman, 1962). The PFR faces axonemal microtubules doublets 4 to 7 and makes several connections with the axoneme. In particular, a thick fibre connects microtubule doublet 7 to the PFR (Sherwin and Gull, 1989). A further contact located between the PFR and the flagellar membrane towards the cell body has also been identified (Sherwin and Gull, 1989). The flagellum attachment zone (FAZ) is a large macromolecular structure located at the cellular membrane facing the flagellum. It is composed of a filament spanning the cellular and flagellar membranes, a set a four microtubules called the microtubule quartet and a FAZ-associated reticulum. The FAZ filament is not present in regions in which the flagellum is intracellular (*i.e*. the flagellar pocket zone). Studies have shown that it starts after the collar, which delimits the intracellular localisation of the flagellum, and ends at the cell body anterior end. The FAZ interdigitates between subpellicular microtubules which are forming an array below the cellular membrane of *T. brucei*. In mature cells, it is present along the whole interface between the flagellum and the cell body. The FAZ, and more particularly the FAZ filament is viewed as a main connecting element with a strong implication in the transfer of flagellum motility to the cell body.

Important knowledge on the FAZ filament composition has been collected from immunoprecipitation, immunofluorescence and bioinformatics (Hu et al., 2015; McAllaster et al., 2015; Moreira et al., 2017; Morriswood et al., 2013; Rotureau et al., 2014; Sunter et al., 2015; Vaughan et al., 2008; Zhou et al., 2015, 2011). The localisation of known FAZ filament proteins and a putative model of their interaction have been presented (Sunter and Gull, 2016). Proteins FAZ1 to FAZ3, FAZ5, FAZ8 to FAZ10 and CC2D localise along the FAZ filament whereas others proteins such as FAZ4, FAZ6, FAZ7, FAZ12 to FAZ14, TbSAS4 and TOEFAZ1 localise to the distal tip of the filament only. Using fluorescence, it has been shown that FAZ11 mainly localises at the FAZ filament distal tip but also possesses a dim localisation along the FAZ filament. Localisations of proteins FAZ15 to FAZ17 have not been identified yet. It has been proposed that the FAZ filament grows by proximal addition of proteins in either a “*push*” or a “*pull*” treadmill-like mechanism (Sunter and Gull, 2016). In the “*push*” model, the proximal addition of structural elements is thought to push the whole FAZ structure, whereas in the “*pull*” model a distal component, yet to be determined, is present in the flagellum compartment and is thought to pull the whole FAZ structure. The FAZ filament is a large macromolecular complex whose structure has previously been investigated using TEM, but not as extensively as other cytoskeletal elements such as the axoneme or the PFR. Indeed, classical TEM studies can only be used on thin specimens (< 250 nm), which is not compatible with the 10 µm-long FAZ filament that coils around a micrometre-thick cell body in mature cells. Several thinning strategies have been used to circumvent this issue: i) conventional resin sections (Sherwin and Gull, 1989), ii) cryo-sections (Höög et al., 2012) and iii) generation of thin anucleated mutant cells for global observation by cryo-TEM (Sun et al., 2018). The punctuated periodic structure of the FAZ filament has been visualised since the early studies on resin sections of heavy-metal stained cells. It is very tempting to associate this punctuated structure to repeated structures also visible in some fluorescence images. The FAZ filament has been first described as a mostly cytoplasmic filamentous structure (Sherwin and Gull, 1989) and later as extracellular staples (Höög et al., 2012). The fact that the main components visible in TEM are intracellular or extracellular has tremendous implications for the identification of the nature of these densities and for the FAZ assembly and elongation mechanisms. It is not clear if this absence of consensus originates from differences between cell types (procyclics versus bloodstream forms) or from biases inherent to some sample preparation methods (conventional TEM versus cryo-TEM). Most of the resin-embedded works show that the FAZ filament is cytoplasmic (Buisson and Bastin, 2010; Sherwin and Gull, 1989). However, heavy-metal staining could reinforce the contrast of these intracellular structures, potentially leading to an exaggeration of their importance compared to neighbouring elements. The natural contrast of structural elements is preserved in cryo-TEM. This lack of consensus could also be explained by the absence of a systematic approach to study longer portions of the FAZ filament since most structural studies were performed on thin sections in which only a thin part of the FAZ filament could be observed.

Cryo-transmission electron tomography (cryo-TET) consists in the collection of projection images of a cryo-fixed sample tilted inside a transmission electron microscope (Frank, 2006). Projection images are then used to computationally reconstruct the object of interest in 3D. It is the method of choice to study macromolecular assemblies and cell components since it allows nanometric resolution imaging of a sample fixed in a close to native state (Lucic et al., 2013, 2008). Nevertheless, cryo-TET is limited to samples thinner than ∼250 nm because of the strong proportion of inelastic scattering occurring in thicker samples (Aoyama et al., 2008). When the sample is too thick, it has to be thinned down using different means such as cryo-sectioning (Höög et al., 2012). Alternatively, people have used smaller cells such as anucleated *T. brucei* (Sun et al., 2018). Scanning transmission electron microscopy (STEM) is an alternative imaging mode, it is based on the raster scanning of the electron beam that is focused on the sample, the transmitted electrons being collected by detectors (Midgley and Weyland, 2003; Pennycook and Nellist, 2011; Sousa and Leapman, 2012). There is no post-specimen electromagnetic lens in STEM and the image contrast (for biological samples) only depends on amplitude contrast as opposed to TEM that relies on phase contrast. Thanks to these differences, STEM is more prone to image thicker samples (above 250nm) as compared to TEM (Aoyama et al., 2008; Biskupek et al., 2010; Hohmann-Marriott et al., 2009; Sousa and Leapman, 2012; Walther et al., 2018). In material sciences, the STEM probe is focused on a very thin sample using high convergence semi-angles to produce very small probes and achieve very high resolutions. In biology of thick samples, the convergence of the electron beam is reduced and ideally the beam is in parallel mode, at the expense of resolution because the probe can be about 1 nm in diameter. The advantage of reducing the convergence semi-angle is to increase the depth of field to several hundreds of nanometres. By combining cryo-methods and STEM tomography (STET), Wolf *et al*. developed the method of cryo-STET in 2014 (Wolf et al., 2014). Simulations have shown that micrometre-thick samples (“*and beyond*”) could be studied using cryo-STET (Rez et al., 2016). However, up to now, no other groups has developed the method, while the cryo-STET pioneers keep on investigating the ultrastructure of biological specimens (Elbaum, 2018; Wolf et al., 2017). Cryo-STET appears as a very promising approach to study cell components *in situ* in thick samples (Wolf and Elbaum, 2019). In the present work, cryo-STET has been developed and applied to study the structure of the FAZ filament in whole chemically-immobilised and cryo-fixed *T. brucei* bloodstream cells. Cryo-tomographic reconstructions confirm that the FAZ filament is composed of a cytoplasmic array of stick-like structures. Systematic study of the FAZ filament along its length unveils that sticks are heterogeneously distributed. Furthermore, sticks are indirectly associated to neighbouring cytoplasmic microtubules via thin appendages whose length varies depending on the type of associated microtubule (on one side, subpellicular microtubule and one the other side, microtubule of the microtubule quartet). Combining the cryo-STET information with protein structure prediction allowed to address new function to FAZ proteins, leading to a new model for the elongation of the FAZ.

## Material and methods

### Sample preparation

*T. brucei* AnTat 1.1E bloodstream forms were cultivated in HMI-11 medium supplemented with 10% foetal calf serum at 37°C in 5% CO2. Exponential growth-phase cells (2×10^6^ parasites/ml) were fixed with formaldehyde (paraformaldehyde 4% w/w final concentration) directly in the culture medium to preserve the cell integrity. A 5 µl drop of the chemically fixed cell culture was deposited on a glow-discharged Quantifoil 200 mesh R2/2 electron microscopy grid (Quantifoil, Großlöbichau, Germany) pre-coated with a gold bead solution. The gold bead solution was composed of commercial 15 nm gold beads (Aurion) and lab-made gold nanorods of various dimensions (synthesised at Li’s laboratory, Ecole Normale Supérieure Chimie Paris-Tech, Paris, France) mixed in equivalent proportions. The grids were manually blotted using Whatman filter paper and plunge-frozen into liquid ethane at −174°C using a Leica EM-CPC equipment (Leica, Wetzlar, Germany). After freezing, the grids were stored in a liquid nitrogen tank until observation by cryo-electron tomography.

### Scanning transmission electron microscopy setup

Frozen electron microscopy grids were mounted on a Gatan 914 high-tilt cryo-holder (Gatan, Pleasanton, CA, USA). Cryo-STET dataset were collected on JEOL 2200FS 200kV field emission gun hybrid TEM/STEM electron microscope (JEOL, Tokyo, Japan). 3k by 3k images were collected in bright-field mode using an on-axis JEOL STEM detector placed at 60 cm camera length. The voltage of the first extraction anode was reduced to 2.1 kV, generating a 1.2 pA beam current at the sample level. A 40 µm condenser aperture was used. In such conditions, the beam convergence and collection semi-angles were 9.3 and 6.6 mrad, respectively. The depth of field associated to the 9.3 mrad convergence semi-angle is about 50 nm. The 9.3 mrad convergence semi-angle has been chosen to generate a small probe diameter to allow small pixel sizes without oversampling. Based on simulations, the probe diameter of an electron beam with a 9.3 mrad convergence semi-angle and a 1.2 pA beam current has been estimated to be around 0.15 nm on an uncorrected JEOL 2200FS (Watanabe et al., 2006). In practice, the probe diameter must be greater than this value because of aberrations. Such convergence semi-angle is associated to strong beam broadening, especially in thick samples, which would deteriorate the images quality is the entire beam was collected. Thus, only a portion of the entire beam has been collected, using a 6.6 mrad collection semi-angle. Dwell time was set to between 1 and 3 µs/pixel and magnifications used ranged between 30,000x and 50,000x (corresponding pixel sizes ranged between 2 and 1.3 nm respectively). A summary of the collection conditions can be found in the Supplementary Table 2. The analogue signal of the bright-field STEM detector was digitised to 16 bit values using a Digiscan II ADC (Gatan, Pleasanton, CA, USA).

### Cryo-STET data acquisition

Images and tilt-series were collected in Digital Micrograph, which is the user interface for controlling the Digiscan II. Digital Micrograph offers scripting possibilities to perform specific and redundant tasks in an automated way. Fully-automatic cryo-STET tilt-series were collected using a homemade script developed in Digital Micrograph. The STET acquisition software used here has been presented in details (Trépout, 2019). Briefly, focusing and tracking tasks are performed on a common region that is localised immediately next to the region of interest. This strategy allows to perform low-dose acquisition. Generally, tilt-series were collected between −70° and +70° using 2° tilt increments. The total electron dose received by the sample ranged between 40 and 80 e^−^/Å^2^. Collection conditions varied from one tilt-series to the other. Thus, the collection conditions of all tilt-series are available in the Supplementary Table 2. In practice, the completion of a whole tilt-series acquisition consisting of ∼70 images took ∼90 minutes.

### Image analysis and segmentation

Fiducial-based alignment and weighted back-projection reconstruction of the tilt-series were performed in Etomo (v.4.9.10) (Kremer et al., 1996; Mastronarde and Held, 2017). After reconstruction, 3D volumes were processed using an edge-enhancing noise-reduction anisotropic diffusion filter to enhance ultrastructural details typically using 10 to 20 iterations (Moreno et al., 2018). Exploration of the reconstructed volumes and segmentations were performed in semi-automatic mode using ImageJ (Schneider et al., 2012). Image measurements and statistical analysis were performed using Matlab (The MathWorks Inc., Natick, MA, USA). The interdistance between the sticks has been computed based on plot profiles made on the arrays of the FAZ filament sticks. After smoothing of the data to reduce the noise, the first derivative of the plot profiles was used to identify the centres of the sticks. Interdistance corresponds then to the distance from the centre of a stick to the centre of the next one. One-way ANOVA statistical tests were performed to measure the p-value for the null hypothesis that the means of the groups are equal (Hogg and Ledolter, 1987). Stick heights and widths were measured manually. The height measurements do not take into account the part of the sticks that might be embedded in the cytoplasmic membrane. The movies were generated in Amira (ThermoFisher Scientific, Hillsboro, OR, USA) and ImageJ (Schneider et al., 2012).

### Protein structure prediction and rendering

A set of 8 FAZ filament proteins (FAZ1 to FAZ3, FAZ5, FAZ8 to FAZ10 and CC2D) were submitted to Phyre2 (http://www.sbg.bio.ic.ac.uk/phyre2/html/page.cgi?id=index) for 3D structure prediction. Phyre2 structure prediction is based on protein homology against a fold library (Kelley et al., 2015). At the time of the study, the fold library contained 71843 entries. Intensive modelling mode was used. FAZ10 is giant protein (0.5 MDa) that could not be modelled as a whole because of Phyre2 sequence size limitation. The FAZ10 protein sequence has then been divided into 5 segments of about 170 kDa each. Two consecutive segments shared an overlapping sequence of 85 kDa not to miss any potential domain prediction. FAZ filament protein structures predicted with high confidence and that contained structural domains greater than 10 nm were rendered using ChimeraX (Goddard et al., 2018). High confidence signifies than at least half of the protein sequence has been modelled with more than 90% confidence.

## Results and discussion

### Ultrastructural organisation of T. brucei

Cells were chemically-fixed prior to cryo-fixation to preserve their integrity. Indeed, *T. brucei* are fragile cells, whose membrane can easily disrupt during the blotting and/or plunge-freezing processes. This fragility has been observed on cells that were not chemically-fixed before freezing (Fig. S1). After immobilisation with formaldehyde, cells were deposited on electron microscopy grids, cryo-fixed in liquid ethane and imaged by cryo-STET. *T. brucei* cell cultures are heterogeneous and contain cells at different stages of the cell cycle. In mature cells, the flagellum and the FAZ are fully grown. Because whole cells were used without any cutting, it was possible to determine accurately the maturation state of the cells. This identification would have been made much more complicated if resin- or cryo-sections would have been used since sections can only contain a portion of the cell. This work focuses uniquely on fully mature *T. brucei* bloodstream cells. In cryo-tomograms, *T. brucei* bloodstream cells display the expected morphology and are embedded in amorphous ice (Fig. 1). In some rare cases, the ice forms crystals and these regions are excluded from analyses (Fig. 1, white asterisks). The tomographic reconstruction contains the whole depth of the cell such that the entire nucleus is visible (Fig. 1A-C, N). The strong contrast allows the visualisation of the nucleolus which appears darker than the rest of the nucleus (Fig. 1A-B, white number sign). Furthermore, connections between the inner and the outer membranes of the nuclear envelope reveal the presence of nuclear pore complexes even at this relatively low magnification (Fig. 1B, yellow arrows). Details of the nuclear envelope are discernible in the magnified view (Fig. 1B, insert). The lysosome is detected on several slices of the reconstruction (Fig. 1A-C, L). In the posterior region of the cell body, a small part of the condensed DNA of *T. brucei* single mitochondrion called the kinetoplast (Fig. 1B, K) is visible next to the flagellar pocket (Fig. 1B, FP).

**Figure 1.**
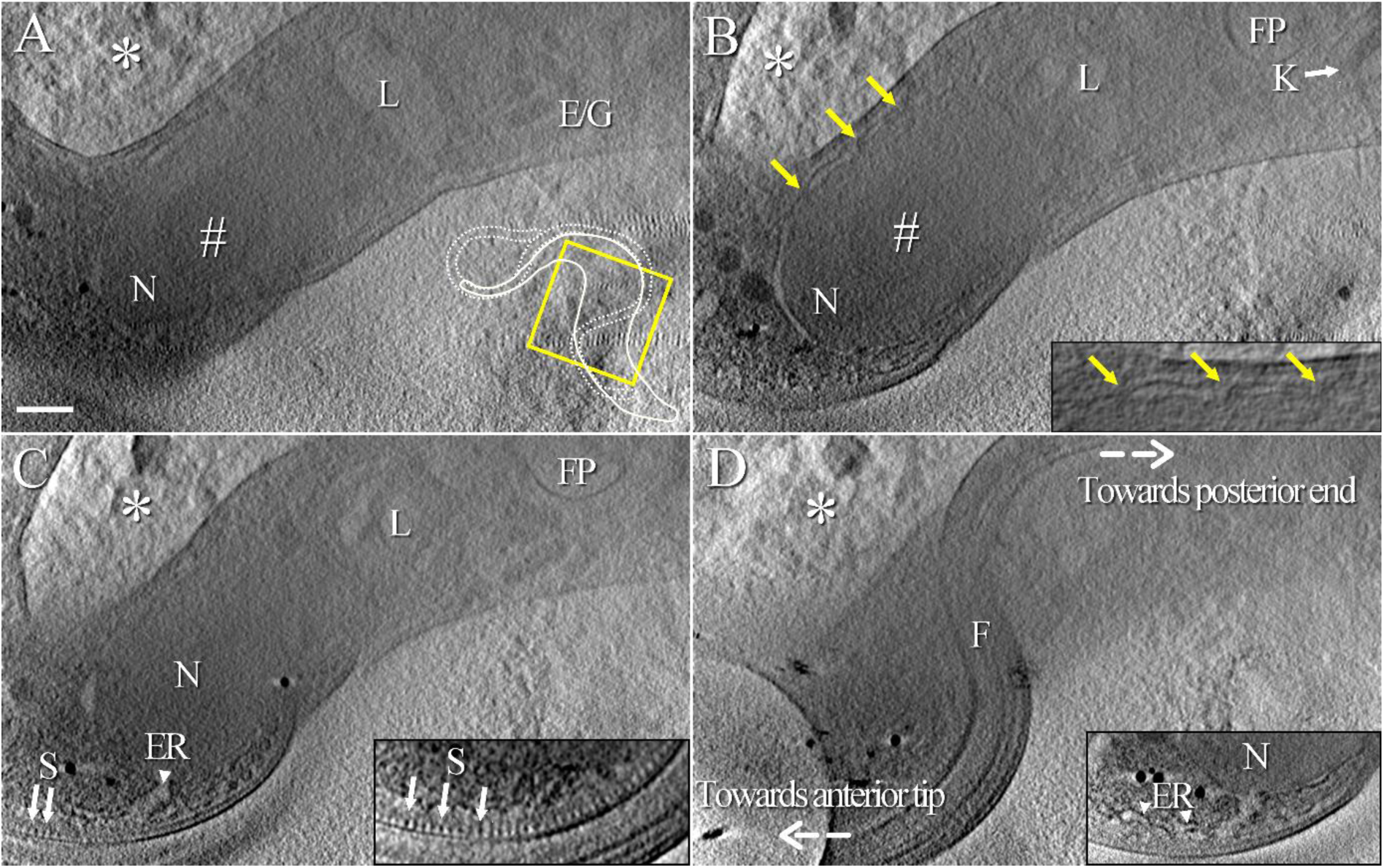
Ultrastructural organisation of a bloodstream *T. brucei* cell observed in cryo-STET. Images are 40 nm-thick slices, spaced by 200 nm, made through a tomographic reconstruction, showing various structural elements found in *T. brucei*. A) Slice passing through the nucleus (N), the nucleolus (#), the lysosome (L) and some endosomes and/or glycosomes (E/G). In the bottom right corner, the yellow square and the small cartoon show which part of the cell is studied in this figure. B) On this second slice, the kinetoplast (K, white arrow), the flagellar pocket (FP) and the location of some nuclear pore complexes (yellow arrows) are visible. The insert is a close-up view of the nuclear envelope. C) Regularly spaced stick-like dark densities (S, arrows) corresponding to the FAZ filament are located next to the FAZ-associated reticulum (ER, arrowhead). The insert is an oriented slice passing through the region of the FAZ filament in which the stick array (S) is visible on a larger scale. D) The last slice shows the flagellum (F) coiled on top of the cell body. The insert is an oriented slice showing the continuity between the outer membrane of the nucleus (N) and the membrane and the lumen of the FAZ-associated reticulum (ER). Directions towards posterior and anterior ends of the cell are indicated with dashed white arrows. The white asterisk in the top left corner of each slice points out at crystalline ice. The whole thickness of this tomogram is about 1.6 µm. The scale bar represents 400 nm.

The FAZ filament appears as a succession of regularly spaced stick-like dark densities found beneath the membrane of the cell body next to the flagellum (Fig. 1C, arrows). Using an oriented virtual tomographic slice, it is possible to better visualise the periodic pattern of these structures (Fig. 1C, insert). The lumen of the FAZ-associated reticulum is observed intracellularly next to the FAZ filament (Fig. 1C, ER). In an oriented virtual tomographic slice, a large part of the FAZ-associated reticulum is visible, its lumen (Fig. 1C) and membrane (Fig. 1D, insert) reaching the intermembrane nuclear space. The flagellum coils along the outer surface of the cell body (Fig. 1D). The visualisation of all these structures is possible since in cryo-STET, images of thick samples can be recorded with sufficient contrast and image content even at high tilts (e.g. greater than ±70°). A movie of this tomogram has been generated to better appreciate the localisation of most of the above-mentioned elements in the cellular context (Movie 1). A movie of the reconstruction is also available (Movie S3).

### Tight contact between cellular and flagellar membranes

We next focused on the space separating the cellular and the flagellar membranes in a cryo-tomogram collected at about 4 µm after the collar of a cell (Fig. 2). In the first slices of the reconstruction, membranes are extremely close to each other (Fig. 2A-B). Then, densities corresponding to the flagellar and the cellular membranes appear slightly separated in the next slices (Fig. 2C-F). When sticks of the FAZ filament are visible, membranes are again pushed against one another (Fig. 2G-I). This proximity can be observed from a side view orientation (Fig. 2J-J’). This membrane close proximity is systematically observed in all collected cryo-tomograms, whatever the location on the flagellum (n=6).

**Figure 2.**
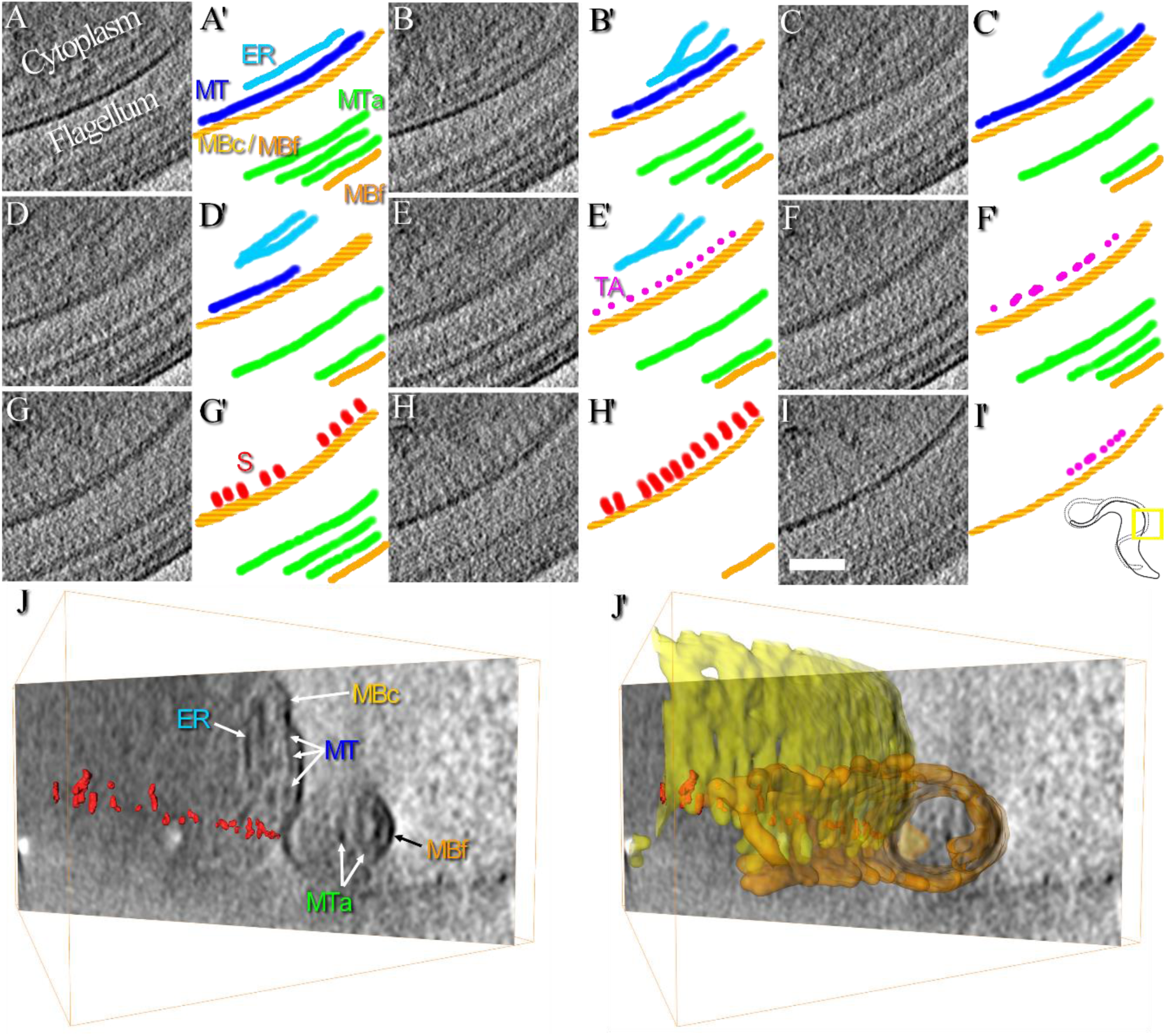
Organisation of the FAZ. This zone corresponds to the area previously displayed in the insert of Figure 1C-D. A-I) Images representing a continuous series of 20 nm-thick consecutive slices made through a tomographic reconstruction showing the structure of the flagellum/cell body interface at about 4 µm after the collar of a cell. A’–I’) Next to each virtual slice, segmentation has been manually realised to highlight the various structures observed. Cellular and flagellum membranes (MBc and MBf, yellow and orange, respectively), the FAZ-associated endoplasmic reticulum (ER, light blue), axonemal microtubules (MTa, green) and a microtubule (MT, dark blue) associated to stick-like structures of the FAZ (S, red) by thin appendages (TA, pink) are highlighted. In the bottom right corner, the yellow square and the small cartoon show which part of the cell is studied in this figure. J) 40 nm-thick oriented slice showing the organisation of the FAZ in a different orientation. All elements presented in panels A-I are indicated here, at the exception of the thin appendages. The microtubule network beneath the cytoplasmic membrane and the proximity between cellular and flagellar membranes are particularly visible in this image. J’) Same image as the one presented in J, showing how the 3D segmented cellular and flagellar membranes are in close contact. The whole thickness of this tomogram is 1.6 µm. The scale bar represents 200 nm.

In previous studies on resin-embedded *T. brucei* procyclic cells, flagellar and cellular membranes are separated by a gap about the size of a microtubule (∼25 nm) (Sherwin and Gull, 1989). To rule out the fact that the difference might arise from cell stage differences, the comparative studies performed on samples from procyclic and bloodstream forms showed that both cell types display a similar gap between flagellar and cellular membranes (Buisson and Bastin, 2010). These gaps were about 10 to 15 nm, smaller than the one observed by Sherwin and Gull (Sherwin and Gull, 1989). It is important to note that the bloodstream form used in Buisson and Bastin is the same strain, cultured in the same laboratory, as the one used in the present work. 10 to 15 nm gaps (*i.e*. 10 to 15 nm) are also observed in bloodstream forms of *T. brucei* (Vickerman, 1962), *T. evansi* (Hiruki, 1987) and *T. congolense* (Vickerman, 1969). Membrane structure is perturbed during sample preparation, especially when dehydration occurs, so further comparison is made with other publications in which cells have been prepared and observed in fully-hydrated state under cryo-conditions. Here, a 30 nm gap is observed between the flagellar and the cellular membranes of *T. brucei* procyclic cells (Höög et al., 2012). The results of the current work agree more with previous resin-embedded study (Buisson and Bastin, 2010) than with cryo-one (Höög et al., 2012).

### The FAZ filament, a mainly cytoplasmic structure made of stick-like densities

Components of the FAZ can be observed in the cryo-tomogram collected 4 µm after the collar of a cell (Fig. 2). The FAZ filament is connected to intracellular microtubules (Sunter and Gull, 2016). These microtubules can be subpellicular microtubules forming the corset or microtubules from the microtubule quartet. It is then expected to find microtubules next to the FAZ filament. In Figure 2, a long structure (Fig. 2, MT, dark blue) with a diameter compatible with that of a microtubule is present beneath the cytoplasmic membrane (Fig. 2, MBc, yellow), in an orientation parallel to the sticks of the FAZ filament (Fig. 2, S, red). Because of its dimension and its localisation, this structure can only be a microtubule. It is not possible to determine if this microtubule belongs to the corset or the microtubule quartet. It is worth noting that a succession of thin and punctuated structures (Fig. 2, TA, pink) is present on both sides of the FAZ filament sticks. The organisation of the FAZ filament, the FAZ-associated ER and a microtubule can also be accessed from a top-view orientation in a reconstruction of the anterior end of another cell (Fig. S2). Sticks of the FAZ filament are 50 nm long and 16 nm wide cytoplasmic entities in contact with the membrane or anchored into it. After denoising of the data using an edge-enhancing noise-reduction anisotropic diffusion filter (Moreno et al., 2018), short densities in the flagellar compartment are sometimes facing cytoplasmic sticks (Fig. 3A, arrowheads).

**Figure 3.**
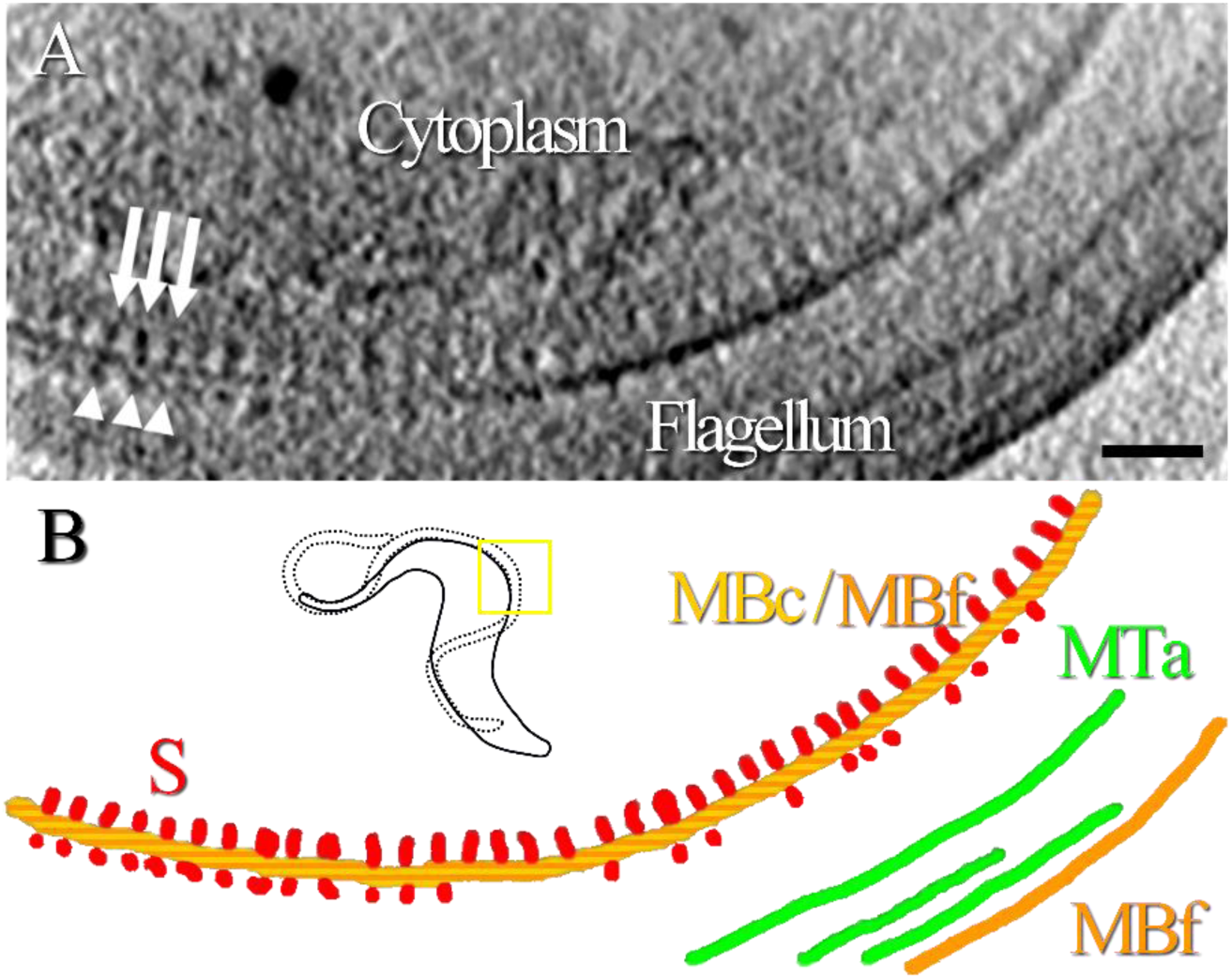
A tightly organised array of intracellular sticks and short flagellar densities. This zone corresponds to the area previously displayed in the insert of Figure 1C. A) Oriented 20 nm-thick slice of a filtered reconstruction (Moreno et al., 2018) in which densities are visible on both sides of the cellular and flagellar membranes. Cytoplasmic sticks of the FAZ filament (white arrows) are longer and more regularly arranged than the flagellar densities facing them (white arrowheads). B) Manual segmentation of the cellular and flagellum membranes (MBc and MBf, yellow and orange, respectively), axonemal microtubules (MTa, green), FAZ cytoplasmic stick-like structures (S, red) and FAZ flagellar short densities (red). The yellow square and the small cartoon show which part of the cell is studied in this figure. The whole thickness of this tomogram is 1.6 µm. The scale bar represents 200 nm.

In previous works, sticks of the FAZ filament are also described as cytoplasmic entities (Buisson and Bastin, 2010) and sometimes, some thin fibrous densities are visible in the flagellar compartment (Sherwin and Gull, 1989). In conventional electron microscopy studies, dehydration of cells together with the use of contrasting agents might increase the visibility of these small structures. In tomography, the sample is not fully tilted inside the electron microscope during the data collection, creating a lack of information in the Fourier space (*i.e*. the missing wedge) which has the effect of blurring the 3D reconstruction in one direction. Moreover, the depth of field is limited when the convergent beam is used in cryo-STET. The missing wedge effect and the low depth of field could also explain why, depending on the orientation of the cell, small flagellar densities are not consistently observed associated to the FAZ filament sticks. Nevertheless, there is no evidence of systematic presence of FAZ filament densities in the flagellum compartment in the present work.

### Sticks are heterogeneously distributed along the FAZ filament

The first cryo-tomograms presented in this work describe the stick organisation at only some positions along the FAZ. To better characterise the stick distribution horizontally along the FAZ filament, several cryo-tomograms were collected at various locations in different uniflagellated cells in a systematic manner to cover most of the FAZ filament. Overall, six cryo-tomograms were collected, each representing about 2 to 3 µm-long portions of FAZ filament. Areas of interest are located at i) the exit of the flagellar pocket, ii) about 4 µm after the collar, iii) about 7 µm after the collar and iv) at the distal end of the FAZ filament (Fig. 4A). To better describe the most proximal and the most distal locations, two tomograms of each zone were collected. A table summarises the position of each tomogram and the figure(s) in which they are displayed (Suppl. Table 2). Based on the analysis of two different cells, no FAZ filament sticks are observed at the proximal region of the flagellum (*i.e*. from the collar up to the first micron of the axoneme) even though the FAZ-associated reticulum is visible (Fig. S3). As observed above, at about 4 µm after the collar the sticks are present and form the regular array of the FAZ filament (Fig. 1-3). On the tomogram collected at about 7 µm after the collar, the curvature of the flagellum is less pronounced and the sticks form an almost straight array (Fig. 5 and Fig. S4). Sticks were previously observed in a top-view orientation at the anterior tip of a cell (Fig. S2). A second tomogram collected at the anterior end of another cell contains side-view orientation of sticks, confirming their presence at the most distal part of the FAZ (Fig. S5).

**Figure 4.**
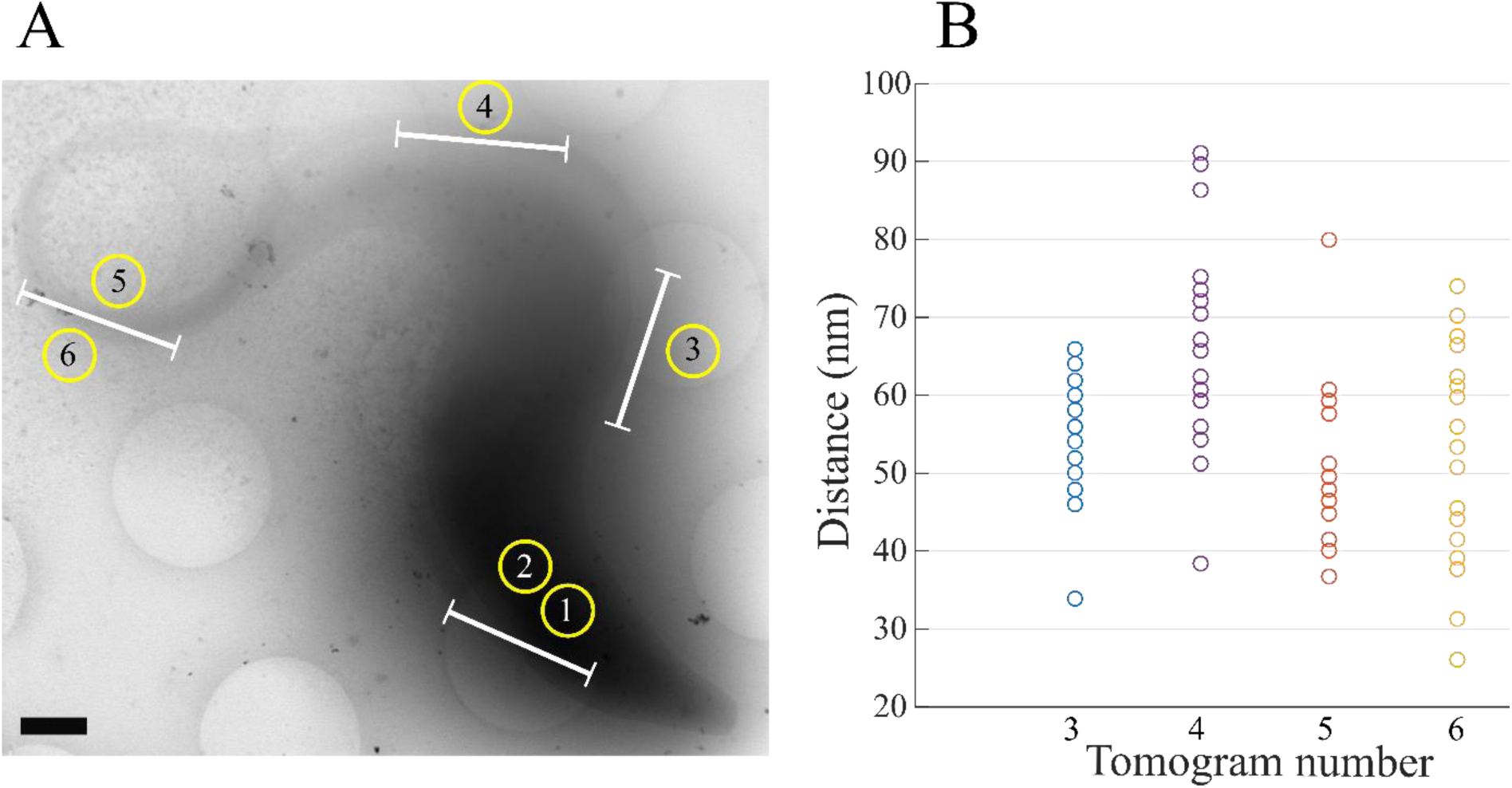
Localisation of investigated FAZ filament portions and measurement of the stick interdistance. Overall six cryo-tomograms were collected to search for the presence of sticks along the FAZ. The distance between two consecutive sticks is measured on the four cryo-tomograms in which sticks are observed. A) Cryo-STEM picture of a *T. brucei* bloodstream cell given as an example to show the positions where the six cryo-tomograms have been collected. Note that cryo-tomograms were collected on different cells. The FAZ portions analysed in each cryo-tomogram are represented by white bars. B) Plot showing the distribution of the stick interdistance for each tomogram in which sticks are observed. The number below each column represents the tomogram number as used in A. Scale bar is 800 nm.

**Figure 5.**
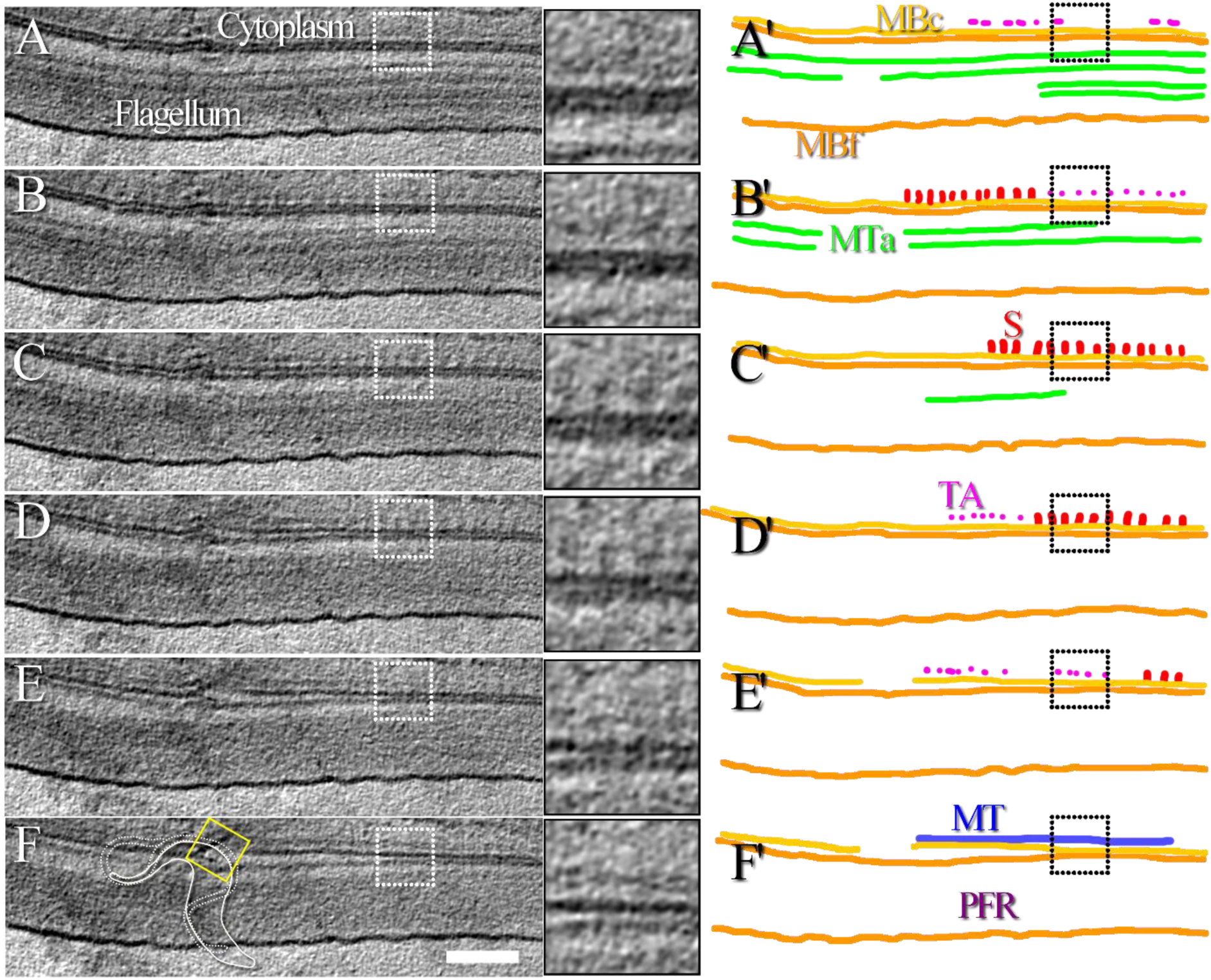
Thin appendages are present between the microtubules and the FAZ filament sticks. A-F) Continuous series of 16 nm-thick consecutive slices made through a tomographic reconstruction showing the structure of the FAZ filament at about 7 µm after the collar of a cell. Each insert represents a magnified view of the original image. The location of the insert is indicated by a dotted white square. In the bottom left corner of image F, the yellow square and the small cartoon show which part of the cell is studied in this figure. A movie of the reconstruction is available (Movie S4). A’-F’) Segmentation highlighting the various structures observed in A-F. The cellular and flagellum membranes (MBc and MBf, yellow and orange, respectively), the paraflagellar rod (PFR), the axonemal microtubules (MTa, green) and the microtubule (MT, blue) separated from stick-like structures of the FAZ (S, red) by thin appendages (TA, pink) are highlighted. The position of the inserts is indicated by the dotted black square. The whole thickness of this tomogram is 0.7 µm. The scale bar represents 250 nm.

In the literature, regularly arranged densities of the FAZ filament observed in electron microscopy are implicitly thought to correspond to FAZ filament proteins which have been detected by immunofluorescence. This association is particularly relevant for FAZ filament proteins that display a punctuated pattern in fluorescence. Following this idea, the absence of sticks at proximal locations as observed in the present study does not agree with immunofluorescence data in which most of the FAZ filament proteins were found to be present at equivalent proximal locations (Kohl et al., 1999; Moreira et al., 2017; Sunter et al., 2015; Vaughan et al., 2008). However, in the work of Moreira *et al*., the signal of FAZ10 is weaker than that of FAZ1 at proximal locations (Moreira et al., 2017). Since FAZ10 is a giant protein (0.5 MDa) it is likely to have a significant contribution in the structure observed in electron microscopy. Its potential relative low abundance at most proximal regions of the FAZ filament might explain why sticks are not observed in the STEM images the present study. Based on this hypothesis, improvement of image quality, either using higher magnification images or 3D reconstruction software taking into account the convergent shape of the beam such as Ettention (Dahmen et al., 2016) or even deconvolution algorithms, might help identifying small protein complexes that would not contain FAZ10. Furthermore, since most of the immunofluorescence works were made on procyclic cells, it might also indicate differences in *T. brucei* cell stages that could be settled on with additional structural and molecular comparative studies.

To further analyse the organisation of the FAZ filament, systematic measurement of the distance between two consecutive sticks is performed (Fig. 4B). The overall mean distance is 67.3±15.7 nm (n=95, including all tomograms). The closest mean distance is observed at the distal end of the FAZ filament (58.4±16.7 nm, n=23) whereas the largest one is measured at about 7 µm after the collar (80.5±16.4 nm, n=20). One-way analysis of variance (ANOVA) shows that measurements are statistically different between these locations on the flagella (p-values ≤ 0.0030) indicating that FAZ filament sticks are not homogenously distributed. ANOVA also shows that the two measurements made at distal ends of FAZ filaments are not statistically different (p-value = 0.7515). These results are in agreement with a heterogeneous horizontal organisation of the FAZ filament sticks. Measurement mean and standard deviation values as well as statistical test results are available as supplementary information (Fig. S6).

Trypanosomes swim forward with the tip of the flagellum leading (Baron et al., 2007; Langousis and Hill, 2014; Walker, 1961). This is due to the fact that beating is initiated at the tip of the flagellum, the waveform being transmitted to the base of the flagellum. It makes sense that FAZ filament sticks are present in high density at the distal tip of the FAZ filament to efficiently attach the flagellum to the cell body during flagellum formation and in mature cells. The observation of FAZ filament sticks at the cell anterior tip is in agreement with the distal localisations of FAZ4, FAZ6, FAZ7, FAZ11 to FAZ14, TbSAS4 and TOEFAZ1 proteins (Hu et al., 2015; McAllaster et al., 2015; Sunter et al., 2015).

### Thin appendages are found between the FAZ filament and microtubules

As observed above in the cryo-tomogram collected about 4 µm after the collar, thin appendages are present between sticks of the FAZ filament and microtubules (Fig. 2, pink). Magnified views of the sticks and appendages are available as supplementary (Fig. S7). These thin appendages are not in contact with the cytoplasmic membrane and their diameter is too small to correspond to microtubules. Because of their localisation and small diameter, these appendages are thought to represent the connections between the FAZ filament and the surrounding microtubules (corset ones and microtubule quartet ones) described in the literature (Sunter and Gull, 2016). To verify that appendages are present along the FAZ filament, all cryo-tomograms are investigated, including the one collected about 7 µm after the collar (Fig. 5). As in Figure 2, a structure whose dimension and localisation allow to identify it as a microtubule is present beneath the cytoplasmic membrane and parallel to the FAZ filament sticks (Fig. 5, MT, blue). Thin appendages (Fig. 5, TA, pink) are visible on both sides of the sticks (Fig. 5, S, red). More slices of this 3D reconstruction are available as supplementary (Fig. S4). Additional images show that the closest microtubule is not at a fixed distance of the sticks depending on which side of the sticks this microtubule is located. By counting the number of slices separating the FAZ filament sticks from the microtubules, it is possible to know the distance between them. The distance on one side is about 50 nm whereas the distance on the other side is about 30 nm.

In the side-view cryo-tomogram collected at the anterior end of a *T. brucei* cell, FAZ-associated reticulum and microtubules are difficulty identified (Fig. S5). Indeed, the FAZ organisation seems different from what has been observed before (Fig. 2). The FAZ organisation could be modified because of a different molecular composition (some microtubules might not reach the anterior tip of the cell) or because of the steric hindrance imposed by the very thin diameter at the cell anterior end (about 150 nm). In previously observed cryo-tomograms, the cell diameter was sufficiently important to accommodate all the components of the entire FAZ. However, when the cell diameter becomes very small, the FAZ might have to organise differently, most probably decorating the whole circumference of the cell, explaining why it is difficult to visualise all components. Nevertheless, thin appendages are present and clearly visible until the anterior tip of the cell body. Because of the small diameter of the cell body in this reconstruction, it is not possible to comment on the distance separating sticks and microtubules.

Based on the observation of three cryo-tomograms, representing over 6 µm of FAZ filament, thin appendages are consistently observed next to the sticks. These appendages represent the microtubule quartet microtubule to FAZ filament domain connection and the FAZ filament domain to subpellicular microtubule connection previously described (Sunter and Gull, 2016). Moreover, since their length varies between ∼30 to ∼50 nm depending on the side of the sticks they locate, this analysis is in favour of the existence of two connections of different nature, yet to be acknowledged. More resolute and detailed analysis is necessary to better describe these connections between FAZ filament sticks and microtubules.

### Towards an identification of the stick nature and function

As mentioned above, the literature implicitly associates the regularly arranged densities of the FAZ filament observed in electron microscopy to the presence of FAZ filament proteins detected by immunofluorescence. To ascertain the identity of the proteins constituting the FAZ filament stick, a comparison is attempted between the structures observed in cryo-STET and predicted structures of FAZ filament proteins. To this purpose, manual measurements were carried out to better describe the stick structure. Sticks are important structures, their average width and height are 16.5 ±4.9 nm and 49.8.2 ±11.7 nm (n=56), respectively (Fig. S8). Current resolution does not allow to comment further on the cylindrical shape or the hollowness of the sticks. However, based on statistical analysis, it is possible that sticks do not have the same dimensions depending on their location on the FAZ filament, sticks being potentially thinner at proximal regions of the FAZ filament (Fig. S8).

Proteins whose localisation (*i.e*. on the FAZ filament) and dimensions (*i.e*. large enough to constitute the sticks observed in cryo-STET) could be compatible are examined. Based on the localisation, these proteins are FAZ1 to FAZ3, FAZ5, FAZ8 to FAZ10 and CC2D (Moreira et al., 2017; Sunter et al., 2015; Vaughan et al., 2008; Zhou et al., 2015, 2011). 3D structure prediction based on protein homology was carried out using Phyre2 (Kelley et al., 2015). Overall, six protein structures are predicted with high confidence (*i.e*. above 50% of the sequence modelled with more than 90% confidence) (Fig. S9). The predicted structures of FAZ1, FAZ2, FAZ8, FAZ9, FAZ10 and CC2D include 10 nm-long (or more) domains mostly made of α-helices, fitting the dimensions of the FAZ filament sticks. More interestingly, domains structurally relevant with desmosome homology are predicted, among which dynein stalk and motor, kinesin stalk, desmoplakin and plakoglobin. The list of predicted relevant domains is available as supplementary (Fig. S10).

A kinesin domain was found in FAZ7, which is present at the distal end of the FAZ filament (Sunter et al., 2015). Subpellicular microtubules have the right polarity for dynein motors to reach the distal end of the cell (Robinson et al., 1995). It is tempting to hypothesise that subpellicular microtubules are used as rails to guide and to extend the FAZ intracellularly. In the present study, the predicted presence of other dynein motor domains in FAZ1, FAZ2 and FAZ10 reinforces the possibility for such mechanism (Fig. S10). Predicted homologies with kinesin and dynein stalk structures concurs with the hypothesis of functional dynein in FAZ filament proteins.

In the literature, the morphological resemblance between FAZ and desmosomes led to the search of proteins with compatible desmosomal structure or function. Bioinformatics analysis on whole *T. brucei* genome identified an armadillo repeat domain similar to that of desmosome proteins in FAZ9 (Sunter et al., 2015). In the present work, Phyre2 also predicted the presence of an armadillo repeat/plakoglobin domain in FAZ9 but also predicted a desmoplakin domain in FAZ1 and FAZ10 (Fig. S10). Most interestingly, FAZ1 and FAZ9 were previously described as potential partners, in full agreement with a desmosome-like structure of the FAZ (Sunter and Gull, 2016).

The potential existence of such domains in FAZ proteins brings more material to elaborate the homology with desmosomes. The prediction of a desmoplakin domain and a dynein motor one in FAZ1 and FAZ10 would place the latter between cytoskeletal elements composed by subpellicular microtubules and the other FAZ proteins. More precisely, the protein FAZ9 and its predicted plakoglobin domain would be the most favourable partner of FAZ1 and FAZ10 (Fig. 6).

**Figure 6.**
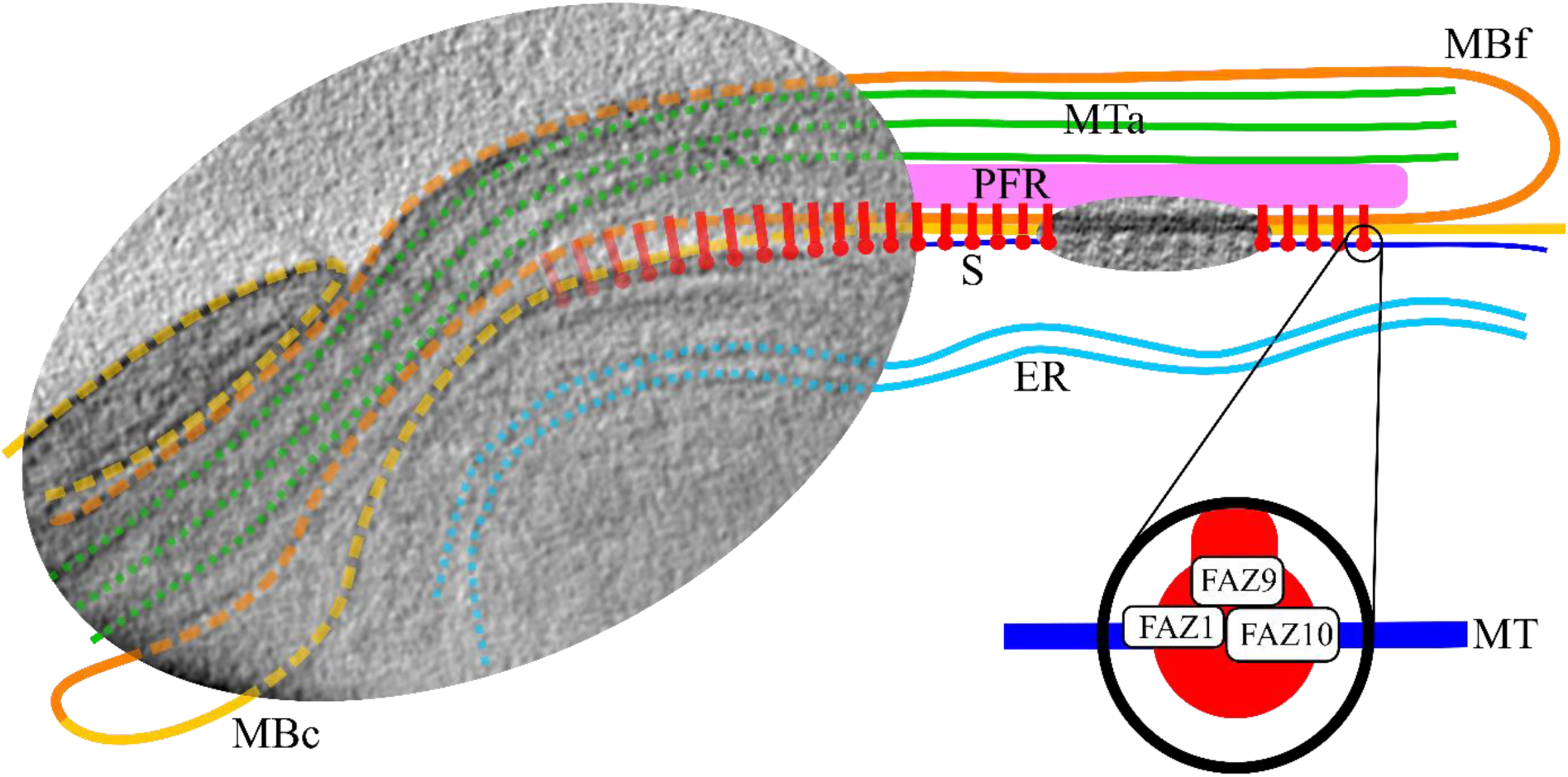
“*Alternative pull*” model of the FAZ elongation. The model is scaled on top of cryo-STET images. Several structures are segmented: cellular and flagellar membranes (MBc and MBf, in yellow and orange, respectively), the FAZ-associated reticulum (ER, light blue), the paraflagellar rod (PFR, pink), axonemal microtubules (MTa, green), a subpellicular microtubule (MT, dark blue) and FAZ filament sticks (S, red) which have been represented joining the cellular and flagellar membranes. Some proximal red structures are faded because their presence has not been confirmed by cryo-STET, yet other publications describe them as being present just after the collar. The sticks are regularly placed following the pattern present in the cryo-STET image. The pulling mechanism is schematised in the zoom-in of a stick. Predicted dynein motor domains of FAZ1 and FAZ10 enable connection with the subpellicular microtubule. Desmoplakin domains of FAZ1 and FAZ10 favour the connection with the plakoglobin domain of FAZ9. Connections with other FAZ filament partners allow the transport of the whole FAZ filament.

The corresponding growing model associated to this FAZ protein organisation would be relatively similar to the “*pull*” model (Sunter and Gull, 2016). Nevertheless, whereas the *“pull”* model involves the presence of a putative protein in the flagellar compartment to elongate the FAZ, the “*alternative pull*” model proposed herein only involves proteins already identified. The driving force of the FAZ elongation would originate from the force exerted by the predicted dynein motor domains of FAZ1 and FAZ10 (and perhaps FAZ2) on subpellicular microtubules (Fig. 6). The pulling would be exerted at each FAZ filament stick location, thus generating a global important pulling force.

## Conclusion

The focus of this work is on the *in situ* ultrastructure and organisation of the FAZ filament in whole *T. brucei* cells using cryo-STET. The observation of typical, textbook-type, intracellular structures attests the good preservation of the cell integrity during blotting and plunge-freezing, especially for such thick sample (cells up to 1.5 µm thick were imaged). The fact that cryo-STET allows to capture large fields of view is an advantage to study eukaryotic cells, it gives the capacity to collect a vast and rich amount of 3D structural information. Thanks to a resolution of a few nanometres, it is possible to describe the heterogeneous organisation of the large FAZ filament, while still being able to capture fine details such as the ones of the thin appendages present between the FAZ filament sticks and the neighbouring microtubules. The current study draws a broader 3D cryo-map of the FAZ filament structure, updating what has previously been observed in conventional electron microscopy of thin sample sections.

The “*alternative pull*” model is based on the combination of i) the confirmed localisation of proteins to the FAZ filament, ii) the dimension of the FAZ filament sticks determined by cryo-STET, iii) the proximity between FAZ filament sticks and cytoplasmic microtubules observed in cryo-STET, iv) the selection of FAZ filament proteins of sizes compatible with the stick dimensions and v) the prediction of structural domains (dynein motor domains and desmosome-like domains). Mutations in the predicted dynein domains of FAZ1 and FAZ10, if they do not perturb the interactions with other FAZ filament proteins, should give a direct evidence of the role of these potential molecular motors in the FAZ filament assembly. Now that important knowledge about FAZ proteins has been gathered and that “*a pattern has emerged linking the RNAi phenotype observed and protein localisation*” (Sunter and Gull, 2016), high resolution structural studies of RNAi phenotypes could extend our understanding of *T. brucei* morphogenesis. Partially-detached flagella phenotype observed in FAZ1^RNAi^ and FAZ5^RNAi^ cell lines (Sunter et al., 2015) are characterised by a mixture of mature and incomplete FAZ structures. A direct structural comparison of these two states would certainly help understanding the complex FAZ organisation. One of the main challenges would be to produce these high resolution maps in a whole cellular environment. Cryo-focused ion beam associated to cryo-TET would most certainly be able to produce such high resolution maps (Schaffer et al., 2015).

Regarding the thickness limitation, this work confirms simulations that stated “*micron thicknesses and beyond*” can be addressed in cryo-STET (Rez et al., 2016). These simulations were made with specific data collection parameters (low convergence semi-angle, nanometric probe size, large collection angle). In the present work, the data collection parameters (relatively high convergence semi-angle, sub-nanometric probe size, limited collection angle) are not qualified as optimal as described in other publications (Rez et al., 2016; Wolf and Elbaum, 2019). This is because of microscope limitations (2 condenser-lens system) that are expected to be circumvented with some tuning of the lens currents in the future. The impact of a 9.3 mrad convergence semi-angle is that resolution in the sample is better at its top compared to its bottom. This has been clearly experimentally demonstrated in the works of Biskupek and Walther, in which parallel beam and convergent beam are compared (Biskupek et al., 2010; Walther et al., 2018). Nevertheless, it makes no doubt that the contrast and the image content presented in this work are of high quality, especially given the low electron dose used and the very thick samples studied. The details on the axonemal microtubules in some of the supplementary movies attest that. This informs about the very strong potential of evolution of the method and the fact that all technical aspects have not been benchmarked yet.

It is important to note that in parallel beam mode, the probe size is about 1 nm, meaning that resolution-wise parallel beam mode is limited. Parallel beam mode has the advantage of a large depth-of-field for imaging cryo-fixed thick samples, but it has the disadvantage of low resolution compared to what is performed in cryo-TET (the wide beam and no scanning mode). Higher resolutions can be achieved in cryo-STET if the probe size if diminished. However, this has multiple consequences, among which the very low depth of field of convergent beams (e.g. in the present study it is about 50 nm). Low depth of field can be compensated using through-focus images as previously demonstrated in the past (Behan et al., 2009; Dahmen et al., 2016; Hovden et al., 2014, 2011; Trepout et al., 2015). By collecting several images at different focal values and combining the different focal planes, it is possible to recover more depth of field. Blurring in the Z direction can also be reduced using 3D reconstruction algorithms that take into account the convergent geometry of the electron beam (Dahmen et al., 2016). These solutions can compensate (at least partly) the inferiority of convergent beams compared to parallel ones demonstrated in previous works (Biskupek et al., 2010; Walther et al., 2018). It would be very interesting to compare through-focus imaging on a convergent beam 200 kV FEG electron microscope versus parallel beam on a 300 kV FEG one. Cards could be reshuffled and it could turn at the advantage of a 200 kV machine instead of that of an expensive 300 kV FEG electron microscope. Performing through-focus imaging require the collection of several images per tilt-angle, hence increasing the amount of electrons. Thus, through-focus imaging should somewhat be compensated to avoid beam damages. Several strategies based on sparse acquisition exist to efficiently reduce the electron dose in STET but have only been applied to non-cryo samples (Li et al., 2018; Trépout, 2019; Vanrompay et al., 2019). These solutions are compatible with through-focus imaging. Cryo-STET has only been developed in 2014 (Wolf et al., 2014), it is a very new method and there are still many opportunities to contribute to its future development.

**Figure caption for Movie 1. Discovery of the structural organisation of a *T. brucei* bloodstream cell**. The movie presents different structural elements present in the cell. The nuclear envelope (NE), the nucleus (N), the lysosome (L), the endoplasmic reticulum (ER) and the flagellum (F) are first displayed on still images. Then, segmentations of the nuclear outer and inner membranes (Nom and Nim, dark and light blue, respectively), the cytoplasmic and flagellar membranes (MBc and Mbf, yellow and orange, respectively) and finally the FAZ filament sticks (S, red) are displayed in 3D.

## Acknowledgments

The author is greatly indebted to P. Bastin (Institut Pasteur, Paris, France) for making this study possible by giving access to the *T. brucei* material, for critical reading of the manuscript and for fruitful discussions about the biology of *T. brucei*. The author thanks C. Travaillé (Institut Pasteur, Paris, France) for providing *T. brucei* bloodstream samples. J.-P. Michel (Institut Galien Paris-Sud, Châtenay-Malabry, France) is acknowledged for his critical reading of the manuscript. The author acknowledges the Multimodal Imaging Centre at Institut Curie Orsay, for providing access to the cryo-electron microscopy facility.

