## Supplementary Data for "*In situ* structural analysis of the flagellum attachment zone in *Trypanosoma brucei* using cryo-scanning transmission electron tomography"

### Supplementary information

Sylvain Trépout

Institut Curie, Inserm US43, CNRS UMS2016, Université Paris-Sud, Université Paris-Saclay, Centre Universitaire, Bât. 101B-110-111-112, Rue Henri Becquerel, CS 90030, 91401 ORSAY Cedex, FRANCE

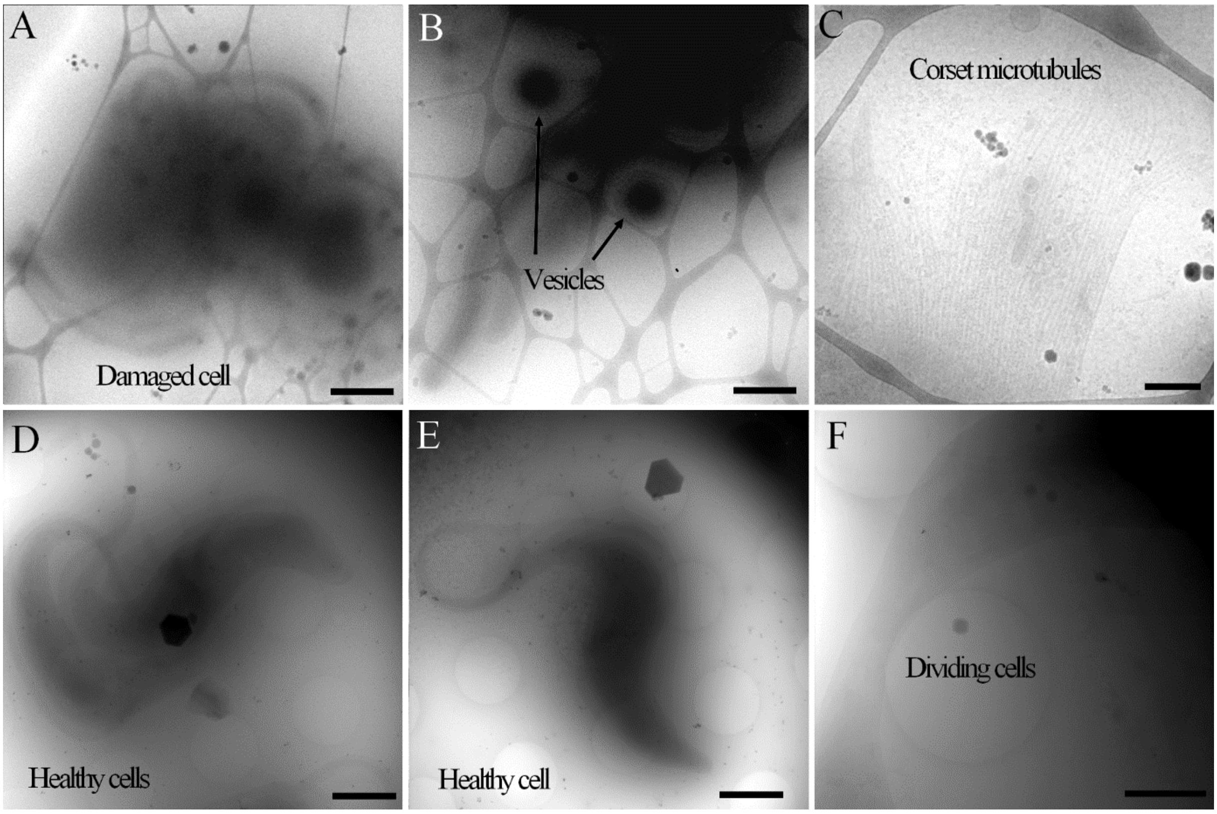

**Supplementary Figure 1. Effects of blotting and plunge-freezing on chemically fixed and unfixed cells.** A-C) Cryo-TET images of unfixed cells, manually blotted using a Whatman filter paper and plunge-frozen in liquid ethane using a Leica EM-CPC on lacey grids. Cells are damaged, they lost their original shape (as observed in light microscopy). Unusual vesicles are present, most probably originating from damaged cytoplasmic membranes. Corset microtubules are also present in the ice, indicating cell rupture. D-F) Cryo-STET images of chemically fixed (4% paraformaldehyde), manually blotted using a Whatman filter paper and plunge-frozen in liquid ethane using a Leica EM-CPC on Quantifoil grids. Cells are healthy and the original shape (as observed in light microscopy) is nicely conserved. The cell membranes are unperturbed, as shown here in a last moment before cell division (F). Please note that the chemical fixation is the parameter that improved the preservation of the cells during blotting. The difference of grid support has a different purpose. The Quantifoil grid only helped improve the gold beads and gold nanorods distribution to better perform the tracking and focusing steps during tilt-series data collection. Scale bars are 2 µm, 800 nm, 200 nm, 2 µm, 2 µm and 1 µm.

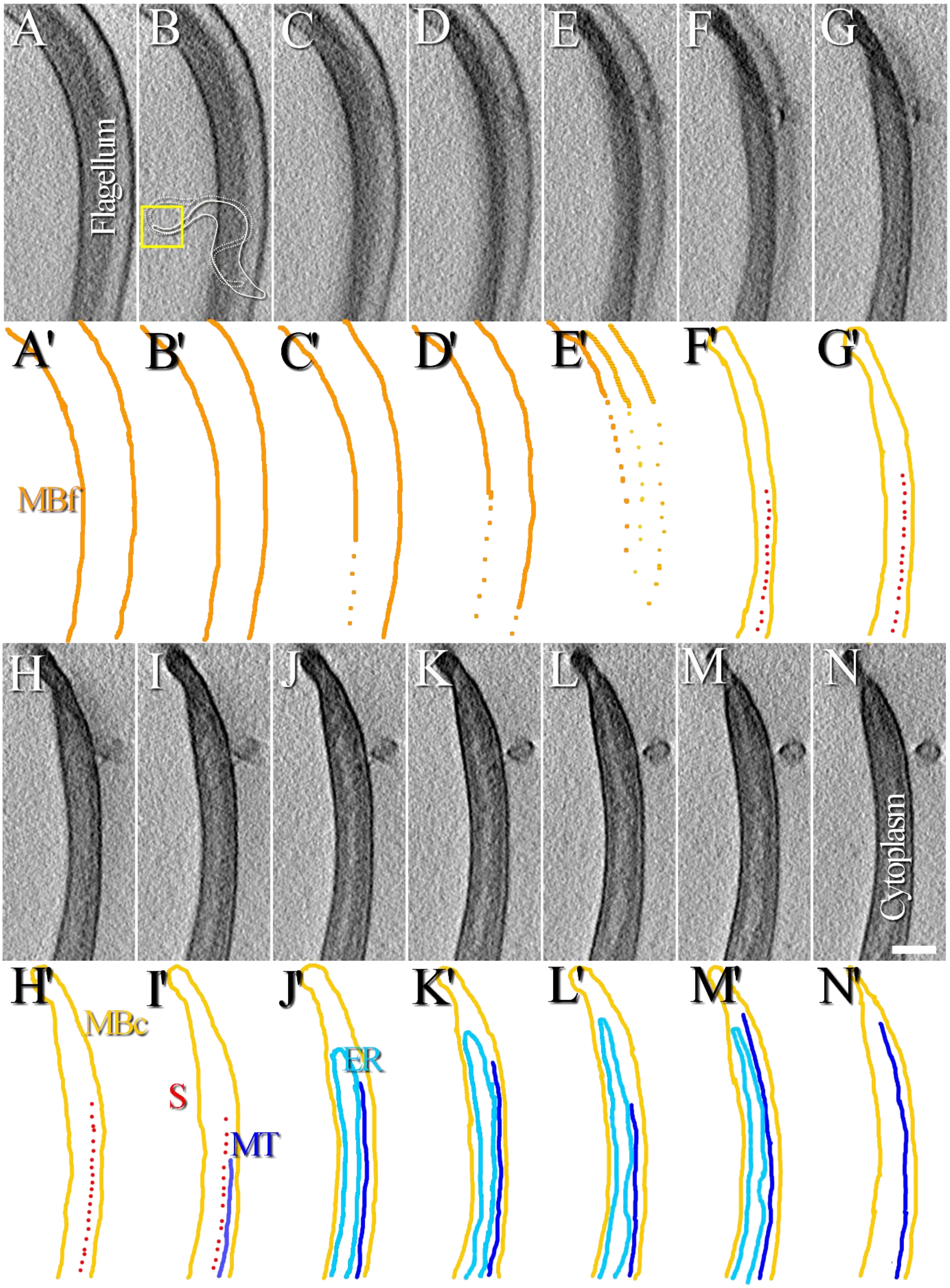

**Supplementary Figure 2. Organisation of the FAZ at the anterior tip of a *T. brucei* cell.** A-N) Images represent a series of 16 nm-thick consecutive slices made through a tomographic reconstruction showing top views of FAZ filament sticks at the cell body/flagellum interface at the anterior tip of a cell. In the image B, the yellow square and the small cartoon show which part of the cell is studied in this figure. Two movies showing the aligned tilt-series and the 3D reconstruction are available as supplementary (Movies S5 & S6). A'–N') Next to each tomographic slice, a segmentation is drawn to highlight various structures observed. Cellular and flagellar membranes (MBc and MBf, yellow and orange, respectively), the FAZ-associated endoplasmic reticulum (ER, light blue), microtubules (MT, dark blue) and stick-like structures of the FAZ filament (S, red) are highlighted. The FAZ-associated ER is visible until the anterior end the cell. The segmentation of the FAZ filament sticks does not reach the cell anterior tip because the cytoplasm crowdedness prevents it. In another reconstruction, sticks do reach the cell anterior tip (Suppl. Fig. 4). Since the thickness of a single slice represents 16 nm, the microtubule segmentation does not correspond to a single microtubule but rather to at least three of them, though it is not possible to resolve them in the reconstruction. The whole thickness of this tomogram is 0.7 µm. The scale bar represents 250 nm.

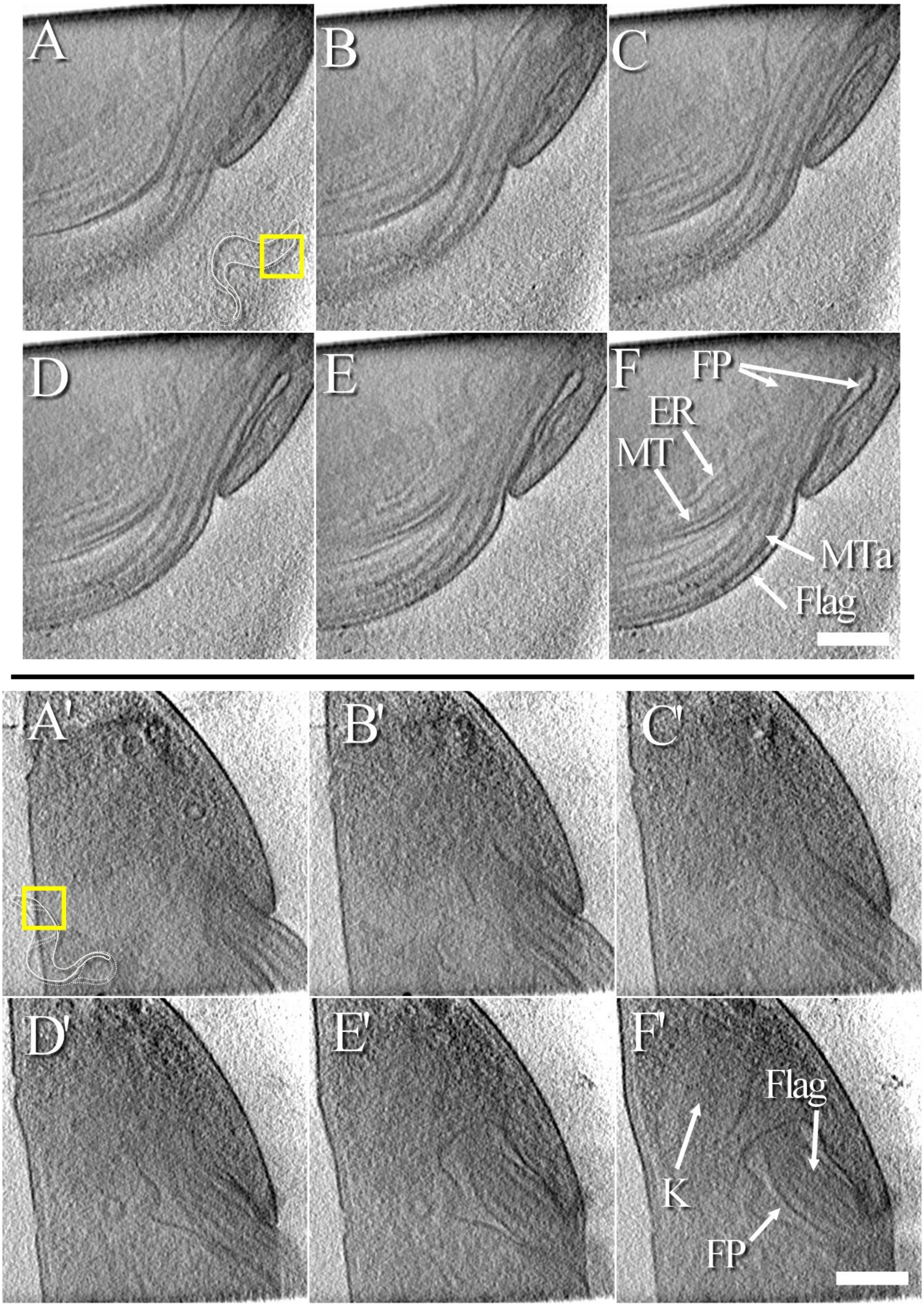

**Supplementary Figure 3. Searching for FAZ filament sticks near the exit of the flagellar pocket.** A-F) Series of 13 nm-thick consecutive slices made through a tomographic reconstruction containing several intracellular structures present near the exit of the flagellar pocket. In the image A, the yellow square and the small cartoon show which part of the cell is studied in this figure. The whole thickness of this tomogram is about 1.3 µm. Two movies showing the aligned tilt-series and the 3D reconstruction are available as supplementary (Movies S1 & S2). A'-F') Series of 13 nm-thick consecutive slices made through a second reconstruction focusing on the flagellar pocket of the cell. In the image A’, the yellow square and the small cartoon show which part of the cell is studied in this figure. The whole thickness of this tomogram is about 1.2 µm. Both reconstructions show the flagellum (Flag) exiting the flagellar pocket (FP). Slice F) The FAZ-associated endoplasmic reticulum (ER), a microtubule (MT) and axonemal microtubules (MTa) are highlighted in the top reconstruction. Slice F’) In the bottom reconstruction, the kinetoplast (K) is visible in a posterior position compared to the flagellar pocket (FP). The scale bars represent 400 nm.

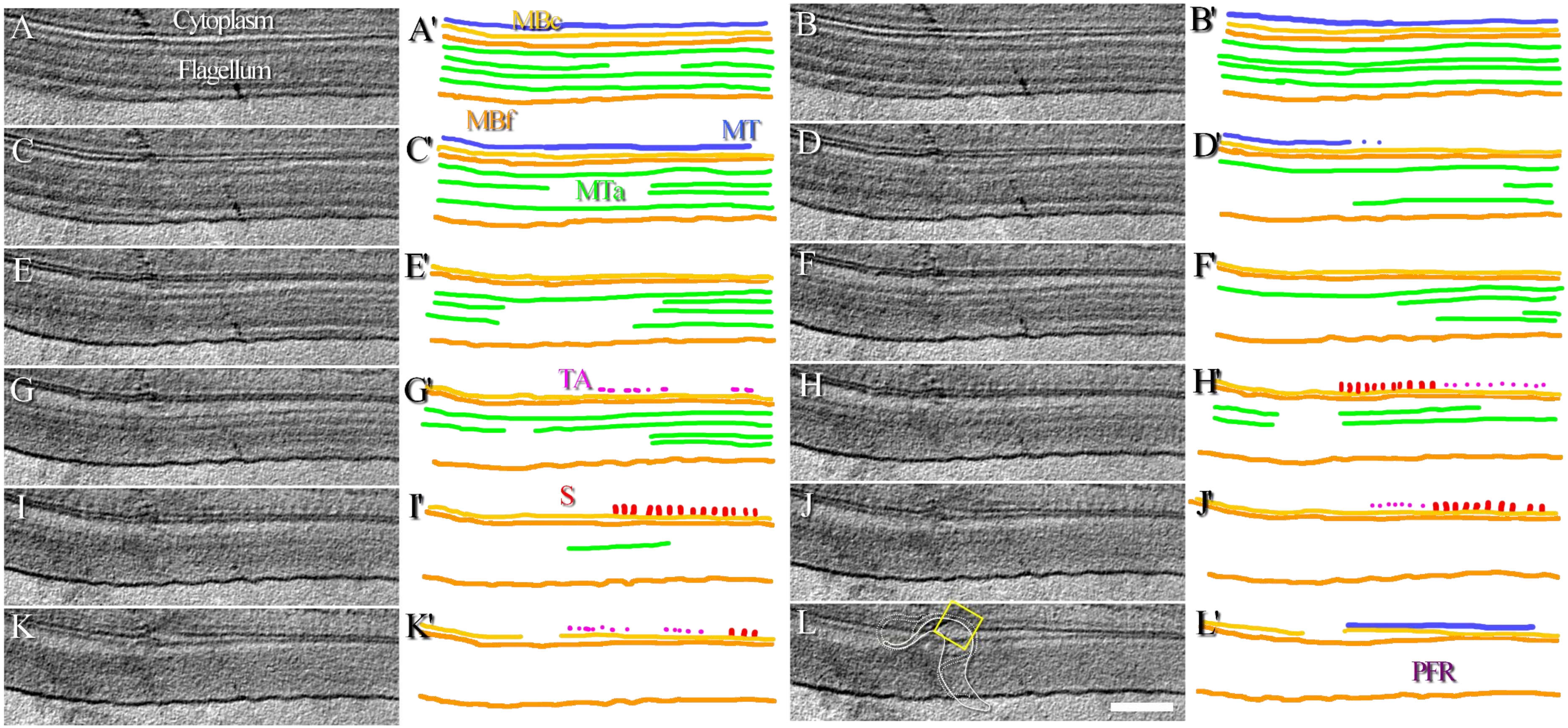

**Supplementary Figure 4. Thin appendages connect microtubules to the FAZ filament sticks.** A-L) Series of 16 nm-thick consecutive slices made through a tomographic reconstruction showing the structure of the FAZ filament at about 7 µm after the collar of a cell. The yellow square and the small cartoon in the image L show which part of a *T. brucei* cell is studied in this figure. A'-L') Segmentation highlighting the various structures observed in A-L. Cellular and flagellum membranes (MBc and MBf, yellow and orange, respectively), the paraflagellar rod (PFR), axonemal microtubules (MTa, green) and cellular microtubules (MT, dark blue) connected to stick-like structures of the FAZ filament (S, red) by thin appendages (TA, pink) are highlighted. The gap between the sticks and the microtubule is wider on the first images (D to H) as compared to the gap on the last images (J to L). This observation informs that the microtubule organisation on both sides of the stick might not be symmetric and that thin appendages could be of different natures. The whole thickness of this tomogram is 0.7 µm. The scale bar represents 250 nm.

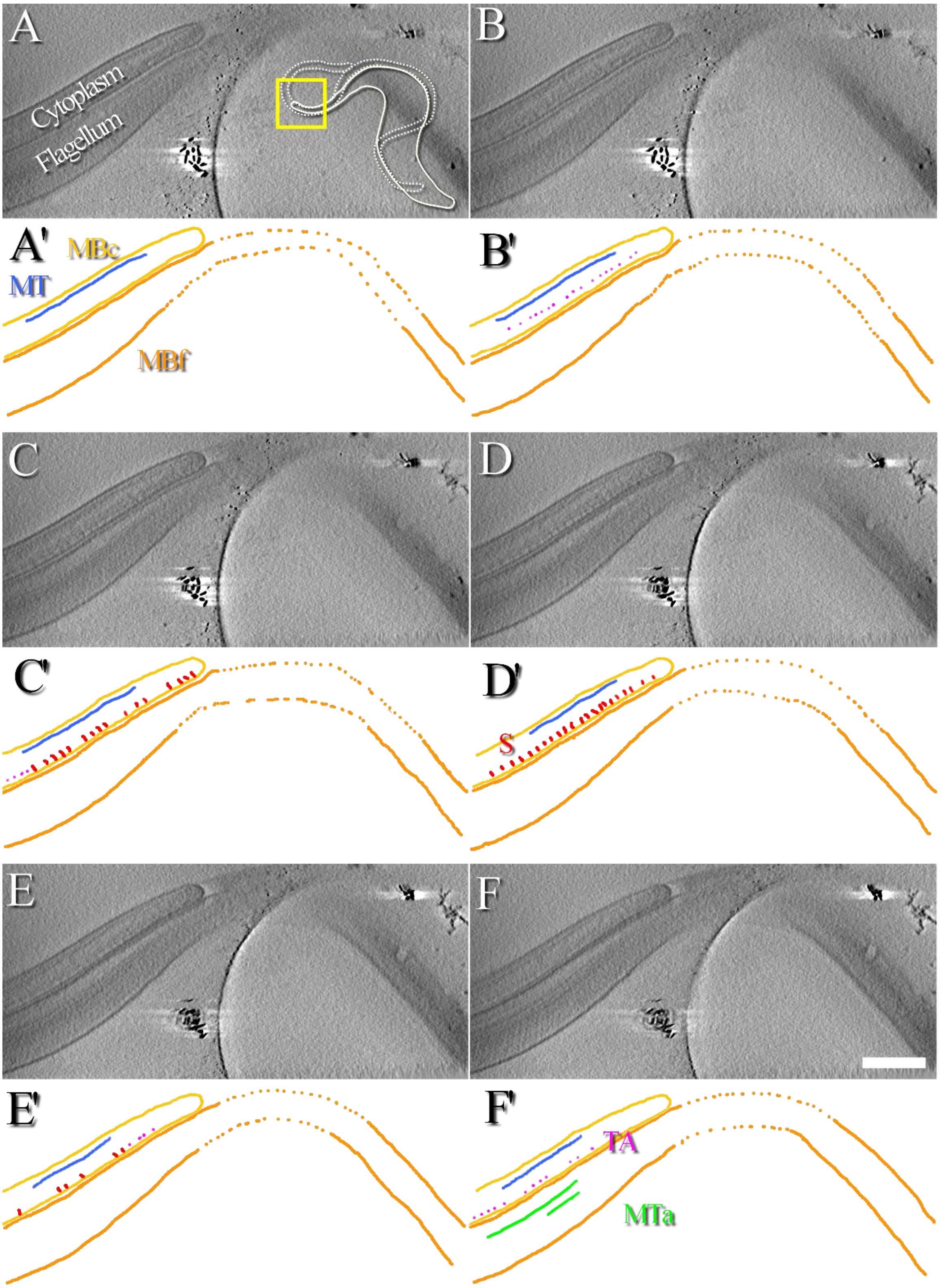

**Supplementary Figure 5. Organisation of the FAZ at the anterior tip of a cell.** A-F) Series of 13 nm-thick consecutive slices made through a tomographic reconstruction showing side views of FAZ filament sticks at the cell body/flagellum interface at the anterior tip of a cell. In the image A, the yellow square and the small cartoon show which part of the *T. brucei* cell is studied in this figure. Two movies showing the aligned tilt-series and the 3D reconstruction are available as supplementary (Movies S7 & S8). A'–F') Next to each virtual slice, various observed structures are highlighted using segmentation. The cellular and flagellar membranes (MBc and MBf, yellow and orange, respectively), axonemal microtubules (MTa, green) and cellular microtubules (MT, blue) connected to stick-like structures of the FAZ (S, red) by thin appendages (TA, pink) are highlighted. The whole thickness of this tomogram is 0.4 µm. The scale bar represents 400 nm.

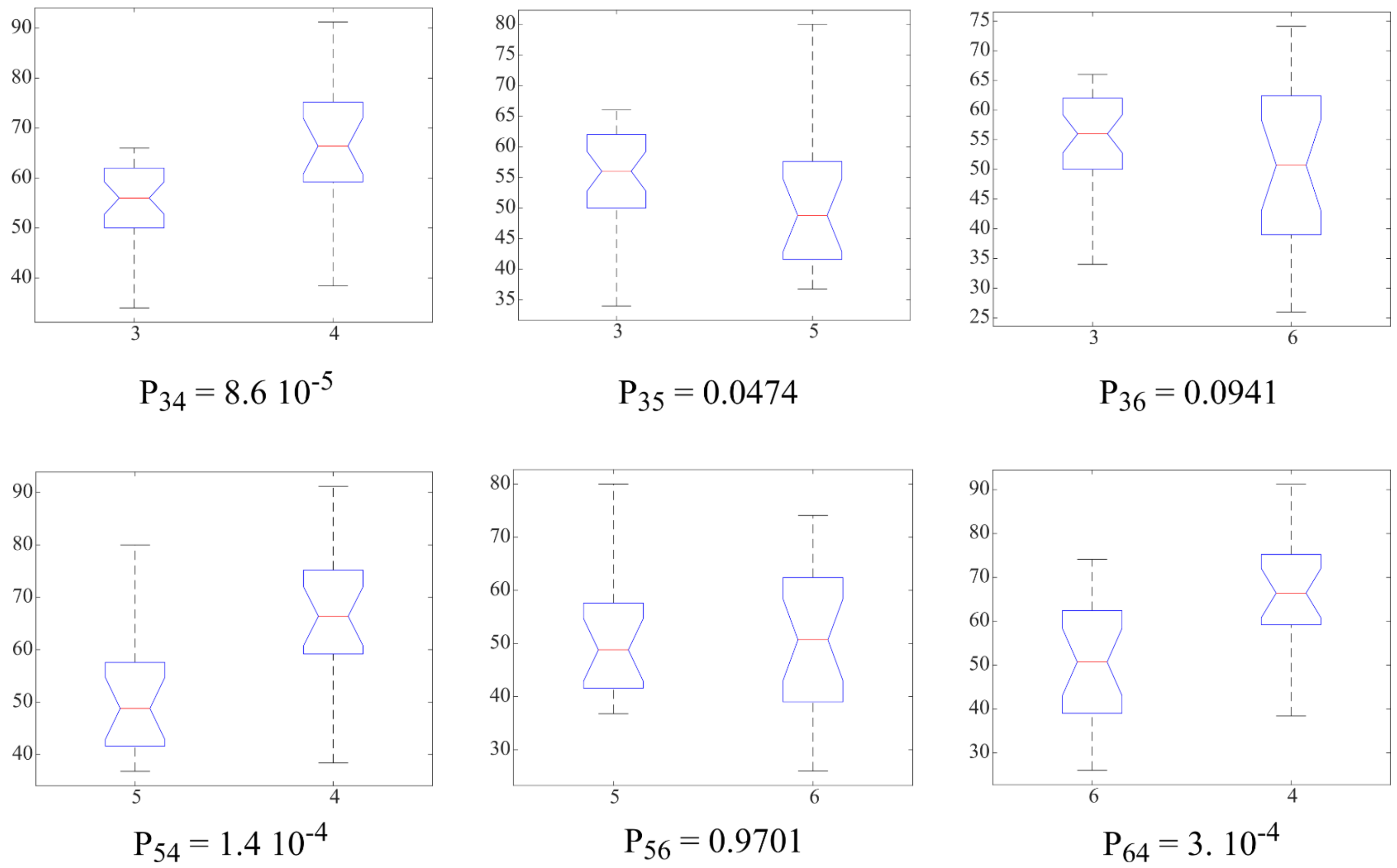

**Supplementary Figure 6. Statistical analysis of the measured distances between FAZ filament sticks.** The notched box plots show the ANOVA results between two different populations. The X axis indicates which population is tested. Populations 3 to 6 correspond to tomogram number as described in Fig. 4A. The Y axis corresponds to the interdistance values expressed in nm. The p-value of the test is displayed below each plot. $P_{34}$ represents the p-value for the test between population 3 and population 4, $P_{35}$ represents the p-value for the test between population 3 and population 5 and so on. Each notched box plot shows the median value (red line), 95% confidence interval of the median (notch), 1$^{st}$ and 3$^{rd}$ quartiles (lower and upper blue lines respectively), interquartile (1$^{st}$ and 3$^{rd}$ quartile interval) and lower and upper whiskers (black lines). The mean and standard deviation values of populations 3 to 6 are 55.6±7.2 nm (n=34), 67.8±13.8 nm (n=20), 50.6±14.5 nm (n=23) and 50.5±10.8 nm (n=18), respectively. Population 4, 7 µm after the collar, is significantly different from any other population. Population 3, 4 µm after the collar, is significantly different from population 5 and there is a 10% risk that it is similar to population 6. However, since the tests indicate that population 5 and 6, both corresponding to anterior end of cells, are statistically not different with a high p-value of 0.9701. Then, it is likely that if more data was available, population 3 would be statistically different form population 6.

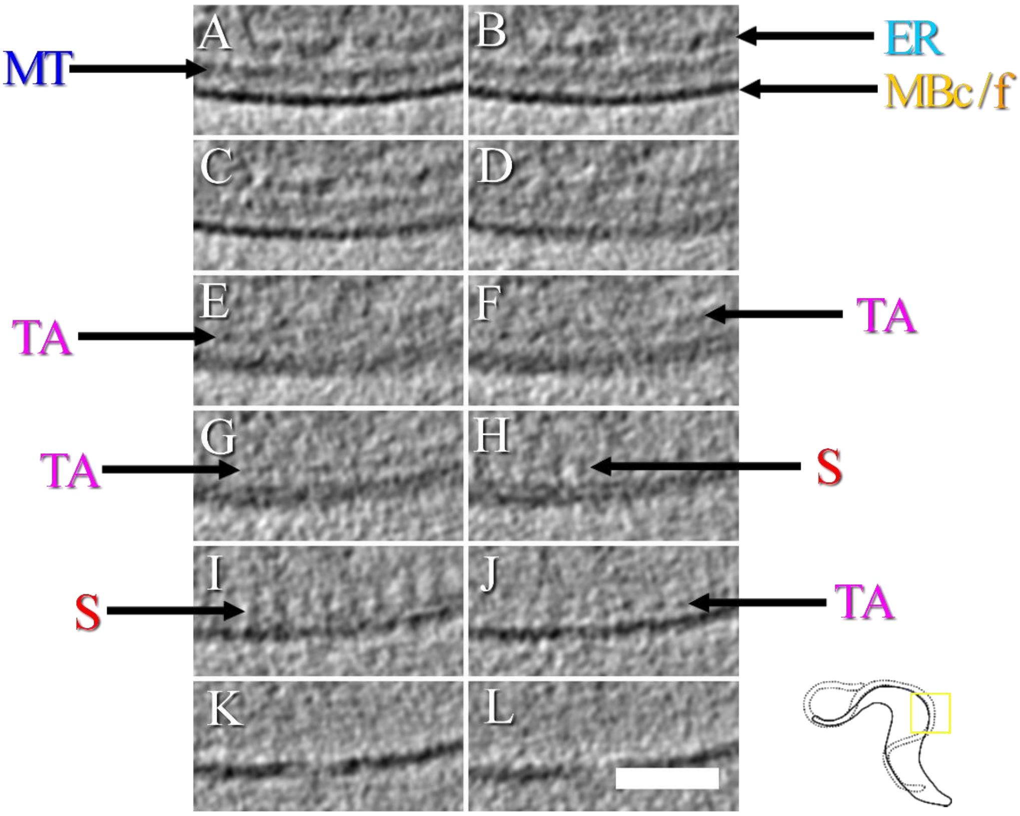

**Supplementary Figure 7. Magnified view of the organisation of the FAZ about 4 µm after the collar of a cell.** This figure presents a magnified view of the zone depicted in Fig. 2. A-L) Series of 20 nm-thick consecutive slices made through a tomographic reconstruction, focusing on the thin appendages present between FAZ filament sticks and a microtubule. The yellow square and the small cartoon in the bottom right part of the figure show which part of the cell is studied. The localisations of both cellular/flagellar membranes (MBc/f), the FAZ-associated endoplasmic reticulum (ER), microtubules (MT), FAZ filament sticks (S) and thin appendages (TA) are indicated. Thin appendages are observed on images E to G, bridging a space of about 80 nm between a microtubule and the sticks, thin appendages are only visible on image J and no microtubule is visible after image J. The whole thickness of this tomogram is 1.6 µm. The scale bar represents 250 nm.

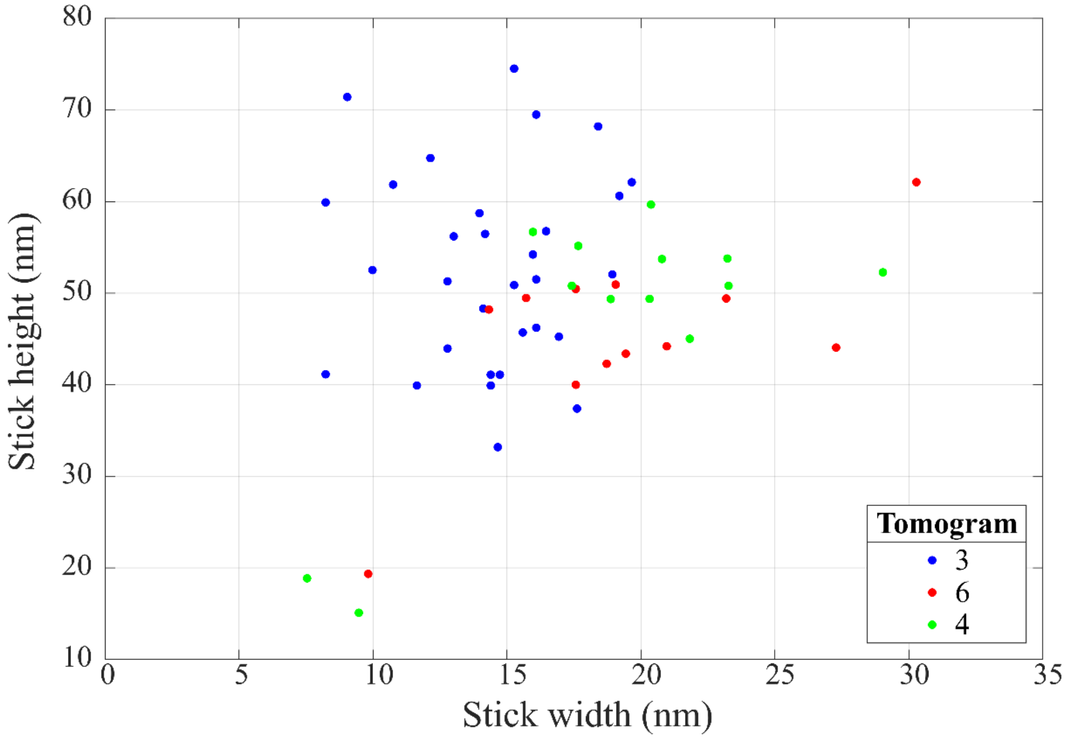

**Supplementary Figure 8. Plot presenting the measured width and height of individual FAZ filament sticks.** Each point corresponds to an individual width/height measurement. Mean width and height are 16.5 ±4.9 nm and 49.8 ±11.7 nm (n=56) respectively. The tomogram number corresponds to the notation indicated in Fig. 4A. Tomogram number 5 is not used for these measurements since it contains top-views of FAZ filament sticks, whose orientation prevents height measurements mainly because of the missing-wedge. Based on ANOVA analysis, the width and height of FAZ filament sticks from tomograms 6 and 4 are statistically not different (p-values = 0.7952 and 0.7379, respectively). The width and height of FAZ filament sticks from tomograms 3 and 6 are statistically different (p-values = 0.004 and 0.0434, respectively). The width of FAZ filament sticks from tomograms 3 and 4 are statistically different (p-value = 0.0014), whereas their height is not (p-value = 0.1394).

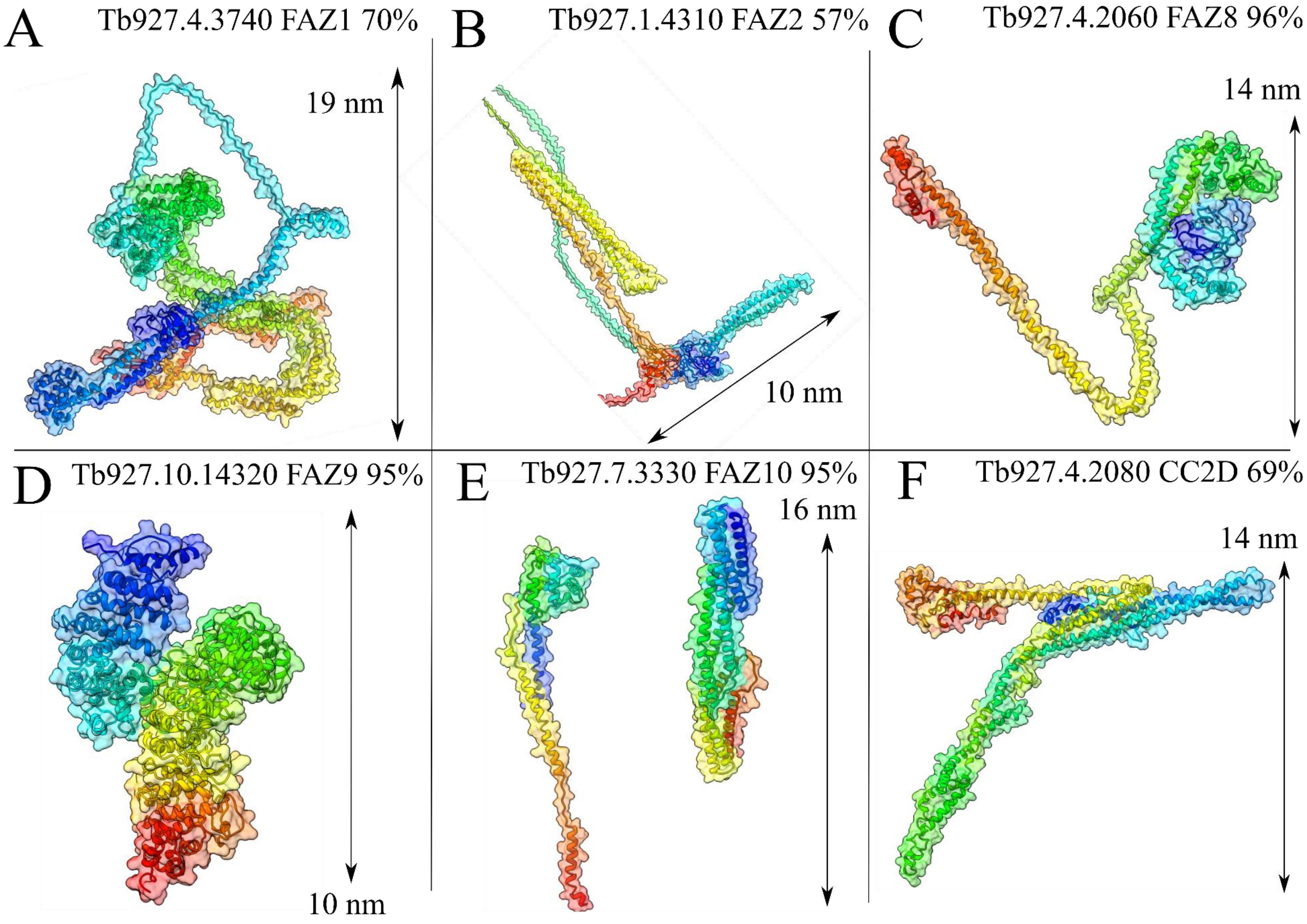
A
Tb927.4.3740 FAZ1 70%
19 nm
B
Tb927.1.4310 FAZ2 57%
10 nm
C
Tb927.4.2060 FAZ8 96%
14 nm
D
Tb927.10.14320 FAZ9 95%
10 nm
E
Tb927.7.3330 FAZ10 95%
16 nm
F
Tb927.4.2080 CC2D 69%
14 nm

**Supplementary Figure 9. Predicted 3D structures of FAZ filament proteins.** Only proteins whose localisation has been identified along the FAZ filament and for which more than 50% of the sequence has been modelled by Phyre2 (Kelley et al., 2015) are presented. The whole modelled sequences of FAZ1, FAZ2, FAZ8 and CC2D are rendered, whereas only selected domains are displayed for FAZ9 and FAZ10. A) 70% of FAZ1 residues are modelled at >90% confidence. B) 57% of FAZ2 sequence is modelled at >90% confidence. C) 96% of FAZ8 residues are modelled at >90% confidence. D) 95% of FAZ9 sequence is modelled at >90% confidence. E) As mentioned in the material and methods section, FAZ10 sequence has been divided into 5 segments of about 170 kDa each with overlapping regions of 85 kDa. From the N-terminal part to the C-terminal one, 91%, 65%, 65%, 71% and 94% of the segment sequences are modelled at >90% confidence respectively. F) 69% of CC2D residues have been modelled at >90% confidence. It should be noted that FAZ11, not represented here because it mainly localises at the distal tip of the FAZ filament, has 99% of its residues modelled at >90% confidence.

| | # | Template | Alignment Coverage | 3D Model | Confidence | % i.d. | Template Information |
|---|---|---|---|---|---|---|---|
| FAZ1 | 3 | c3vkhA_ | Alignment | | 98.9 | 18 | **PDB header:**motor protein **Chain:** A: **PDB Molecule:**dynein heavy chain, cytoplasmic; **PDBTitle:** x-ray structure of a functional full-length dynein motor domain |
| | 40 | c3r6nA_ | Alignment | not modelled | 95.8 | 8 | **PDB header:**cell adhesion **Chain:** A: **PDB Molecule:**desmoplakin; **PDBTitle:** crystal structure of a rigid four spectrin repeat fragment of the2 human desmoplakin plakin domain |
| | 53 | c6ignA_ | Alignment | not modelled | 95.0 | 19 | **PDB header:**motor protein **Chain:** A: **PDB Molecule:**kinesin-1 heavy chain; **PDBTitle:** the crystal structure of kif5b stalk 1 coiled-coil region |
| FAZ2 | 12 | c3wuqA_ | Alignment | | 95.4 | 7 | **PDB header:**motor protein **Chain:** A: **PDB Molecule:**cytoplasmic dynein 1 heavy chain 1; **PDBTitle:** structure of the entire stalk region of the dynein motor domain |
| | 13 | c4rh7A_ | Alignment | | 95.2 | 10 | **PDB header:**motor protein **Chain:** A: **PDB Molecule:**green fluorescent protein/cytoplasmic dynein 2 heavy chain **PDBTitle:** crystal structure of human cytoplasmic dynein 2 motor domain in2 complex with adp.vi |
| FAZ8 | 17 | c3wuqA_ | Alignment | | 91.7 | 9 | **PDB header:**motor protein **Chain:** A: **PDB Molecule:**cytoplasmic dynein 1 heavy chain 1; **PDBTitle:** structure of the entire stalk region of the dynein motor domain |
| FAZ9 | 3 | c3ifqB_ | Alignment | | 100.0 | 22 | **PDB header:**cell adhesion **Chain:** B: **PDB Molecule:**plakoglobin; **PDBTitle:** interction of plakoglobin and beta-catenin with desmosomal2 cadherins |
| FAZ10 | 9 | c3vkhA_ | Alignment | | 97.6 | 19 | **PDB header:**motor protein **Chain:** A: **PDB Molecule:**dynein heavy chain, cytoplasmic; **PDBTitle:** x-ray structure of a functional full-length dynein motor domain |
| | 58 | c6ignA_ | Alignment | not modelled | 94.7 | 17 | **PDB header:**motor protein **Chain:** A: **PDB Molecule:**kinesin-1 heavy chain; **PDBTitle:** the crystal structure of kif5b stalk 1 coiled-coil region |
| | 20 | c3wuqA_ | Alignment | | 97.9 | 9 | **PDB header:**motor protein **Chain:** A: **PDB Molecule:**cytoplasmic dynein 1 heavy chain 1; **PDBTitle:** structure of the entire stalk region of the dynein motor domain |
| | 9 | c5j1iA_ | Alignment | | 98.4 | 15 | **PDB header:**structural protein **Chain:** A: **PDB Molecule:**plectin; **PDBTitle:** structure of the spectrin repeats 7, 8, and 9 of the plakin domain of2 plectin |
| | 25 | c3r6nA_ | Alignment | | 97.6 | 8 | **PDB header:**cell adhesion **Chain:** A: **PDB Molecule:**desmoplakin; **PDBTitle:** crystal structure of a rigid four spectrin repeat fragment of the2 human desmoplakin plakin domain |
| CC2D | 23 | c6ignA_ | Alignment | not modelled | 96.8 | 8 | **PDB header:**motor protein **Chain:** A: **PDB Molecule:**kinesin-1 heavy chain; **PDBTitle:** the crystal structure of kif5b stalk 1 coiled-coil region |
| | 37 | c3wuqA_ | Alignment | | 95.8 | 8 | **PDB header:**motor protein **Chain:** A: **PDB Molecule:**cytoplasmic dynein 1 heavy chain 1; **PDBTitle:** structure of the entire stalk region of the dynein motor domain |

**Supplementary Figure 10. Predicted domains of FAZ filament proteins.** Structural domains of FAZ filament proteins modelled at more than 50% by Phyre2 (Kelley et al., 2015) are listed. The list contains domains structurally relevant for a desmosome-like structure of the FAZ, to which, microtubule-related domains are added. The tables have been extracted directly from Phyre2 hit report results. From left to right, the columns represent: i) the hit number in the Phyre2 result, ii) the name of the corresponding template, iii) the location of the alignment in the context of the whole sequence, iv) a snapshot of the 3D model, v) the overall confidence of the prediction, vi) the percentage of amino-acid sequence homology and vii) a description of the template biological role. FAZ1) N-terminal and C-terminal regions of FAZ1 have homology with dynein motor and kinesin stalk structures whereas a desmoplakin structure is predicted at the centre of the sequence. FAZ2) The whole first half of the sequence is predicted disordered. The second half is predicted to contain dynein stalk and motor domains. FAZ8) A dynein stalk domain is predicted in the second half of the protein sequence. FAZ9) The predicted tertiary structure of the sequence second half is an armadillo repeat of plakoglobin at 100% confidence. FAZ10) The N-terminal region of the protein is predicted to contain a dynein motor domain. Kinesin and dynein stalk domains are predicted in the first half of the protein sequence. Plakin and desmoplakin domains are predicted to be present at the centre of the sequence. CC2D) N-terminal and C-terminal regions are predicted to contain kinesin and dynein stalks domains respectively. It should be noted that FAZ11, not represented here because it mainly localises at the distal tip of the FAZ filament, is predicted to contain stalk regions of kinesin and dynein motors.

| Tomogram 3 (n=34) | Tomogram 4 (n=20) | Tomogram 5 (n=18) | Tomogram 6 (n=23) |
|---|---|---|---|
| 48.0 | 38.4 | 59.2 | 26.0 |
| 56.0 | 59.2 | 57.6 | 39.0 |
| 54.0 | 67.2 | 48.0 | 26.0 |
| 56.0 | 51.2 | 57.6 | 39.0 |
| 46.0 | 72.0 | 51.2 | 31.2 |
| 58.0 | 60.8 | 46.4 | 62.4 |
| 56.0 | 62.4 | 41.6 | 67.6 |
| 66.0 | 65.6 | 80.0 | 67.6 |
| 62.0 | 59.2 | 41.6 | 44.2 |
| 58.0 | 54.4 | 49.6 | 62.4 |
| 56.0 | 60.8 | 49.6 | 55.9 |
| 50.0 | 75.2 | 44.8 | 66.3 |
| 48.0 | 56.0 | 60.8 | 70.2 |
| 66.0 | 86.4 | 60.8 | 45.5 |
| 62.0 | 89.6 | 36.8 | 59.8 |
| 48.0 | 91.2 | 40.0 | 61.1 |
| 66.0 | 73.6 | 36.8 | 37.7 |
| 56.0 | 86.4 | 46.4 | 53.3 |
| 56.0 | 70.4 | | 50.7 |
| 48.0 | 75.2 | | 74.1 |
| 52.0 | | | 37.7 |
| 46.0 | | | 45.5 |
| 56.0 | | | 41.6 |
| 34.0 | | | |
| 56.0 | | | |
| 58.0 | | | |
| 56.0 | | | |
| 60.0 | | | |
| 66.0 | | | |
| 64.0 | | | |
| 62.0 | | | |
| 56.0 | | | |
| 62.0 | | | |
| 46.0 | | | |
| **Mean = 55.6** | **Mean = 67.8** | **Mean = 50.5** | **Mean = 50.6** |
| **Std = 7.2** | **Std = 13.8** | **Std = 10.8** | **Std = 14.5** |

**Supplementary Table 1. Distances measured between two consecutives FAZ filament sticks.** Values expressed in nm. Mean and standard deviation are given at the bottom of each column.

| Tomogram number: | Displayed in: | Collection conditions |
|---|---|---|
| **Tomogram 1 (exit of flagellar pocket)** | **Supplementary Figure 3 (top)<br>Movies S1 & S2** | **Pixel size: 1.3 nm<br>Dwell time: 1.5 μs<br>Beam current: 1.6 pA<br>Angular range: -64° : 2° : 75°<br>Thickness: 1.3 μm<br>Electron dose: 80 $e^-/Å^2$** |
| **Tomogram 2 (exit of flagellar pocket)** | **Supplementary Figure 3 (bottom)** | **Pixel size: 1.3 nm<br>Dwell time: 1.5 μs<br>Beam current: 1.6 pA<br>Angular range: -58° : 2° : 79°<br>Thickness: 1.2 μm<br>Electron dose: 80 $e^-/Å^2$** |
| **Tomogram 3 (about 4 μm after the collar)** | **Figures 1, 2 and 3<br>Supplementary Figure 7<br>Movie 1<br>Movie S3** | **Pixel size: 2 nm<br>Dwell time:3 μs<br>Beam current: 1.2 pA<br>Angular range: -65° : 2° : 78°<br>Thickness: 1.6 μm<br>Electron dose: 78 $e^-/Å^2$** |
| **Tomogram 4 (about 7 μm after the collar)** | **Figure 5<br>Supplementary Figure 4<br>Movie S4** | **Pixel size: 1.6 nm<br>Dwell time: 2 μs<br>Beam current: 1.2 pA<br>Angular range: -76° : 2° : 75°<br>Thickness: 0.7 μm<br>Electron dose: 65 $e^-/Å^2$** |
| **Tomogram 5 (anterior end of the cell) – top views** | **Supplementary Figure 2<br>Movies S5 & S6** | **Pixel size: 1.6 nm<br>Dwell time: 2 μs<br>Beam current: 1.2 pA<br>Angular range: -72° : 2° : 76°<br>Thickness: 0.7 μm<br>Electron dose: 65 $e^-/Å^2$** |
| **Tomogram 6 (anterior end of the cell) – side views** | **Supplementary Figure 5<br>Movies S7 & S8** | **Pixel size: 1.3 nm<br>Dwell time: 1 μs<br>Beam current: 1.2 pA<br>Angular range: -71° : 2° : 67°<br>Thickness: 0.4 μm<br>Electron dose: 40 $e^-/Å^2$** |

**Supplementary Table 2. Details about tomogram displayed in the present work.** The correspondence between tomogram number (*i.e.* as indicated in Fig. 4A) and figure number (*i.e.* in which the tomographic data is displayed) is indicated**.** This table helps associating which tomogram is displayed in which figure, movie, supplementary figure or supplementary movie. The flagellum location at which the tomogram has been collected is also indicated between brackets. Overall, 15 cryo-tomograms have been collected to help understanding the organisation of the FAZ filament.

**Supplementary Movie 1. Aligned tilt-series of tomogram 1.** The white square delimits a zone with a different brightness/contrast to better appreciate the fact that even at high tilts, there is information arriving to the BF STEM detector.

**Supplementary Movie 2. Reconstruction of tomogram 1.** A simple 2x2x2 Gaussian 3D filter has been applied to the original weighted back-projection to reduce the noise of the images and allow sufficiently good h264 compression.

**Supplementary Movie 3. Reconstruction of tomogram 3.** A simple 2x2x2 Gaussian 3D filter has been applied to the original weighted back-projection to reduce the noise of the images and allow sufficiently good h264 compression.

**Supplementary Movie 4. Reconstruction of tomogram 4.** A simple 2x2x2 Gaussian 3D filter has been applied to the original weighted back-projection to reduce the noise of the images and allow sufficiently good h264 compression.

**Supplementary Movie 5. Aligned tilt-series of tomogram 5.** The white square delimits a zone with a different brightness/contrast to better appreciate the fact that even at high tilts, there is information arriving to the BF STEM detector.

**Supplementary Movie 6. Reconstruction of tomogram 5.** A simple 2x2x2 Gaussian 3D filter has been applied to the original weighted back-projection to reduce the noise of the images and allow sufficiently good h264 compression.

**Supplementary Movie 7. Aligned tilt-series of tomogram 6.** Note the particularly good focusing on the gold nanorods at the bottom of the image (just a very few focus jumps).

**Supplementary Movie 8. Reconstruction of tomogram 6.** A simple 2x2x2 Gaussian 3D filter has been applied to the original weighted back-projection to reduce the noise of the images and allow sufficiently good h264 compression.
